# A pathway-informed mutual exclusivity framework to detect genetic interactions in pediatric cancer

**DOI:** 10.1101/2025.07.14.664743

**Authors:** Anastasia Spinou, Richard Gremmen, Jarno Drost, Patrick Kemmeren

## Abstract

**Background:** The exponential increase of sequenced cancer genomes has enabled the in-silico study of genetic interactions in tumors across many cancer types - particularly synthetic lethality, where two gene alterations lead to cell death - and identify new candidate therapeutic targets. This rise is primarily present in adult cancer, while in-silico investigation of genetic interactions remains challenging in pediatric cancer especially in tumors of low incidence. Consequently, this underscores the need for specialized approaches to advance our understanding of genetic interactions in pediatric oncology.

**Methods:** Here, we present the pathway-informed genetic interaction framework (PIGI) that employs mutual exclusivity and co-occurrence testing and leverages biological pathways to infer candidate genetic interactions. Pathways facilitate the detection of hidden biology by grouping genes in functional units, as well as alleviate key confounders of these analyses: pathway epistasis and cancer subtypes - thereby highlighting genes of greater interest.

**Results:** PIGI detected candidate genetic interactions by assessing pathway mutual exclusivity and co-occurrence in two primary pediatric cancer datasets, DKFZ and TARGET. PIGI detected, from high significance pathway relationships, 35 mutually exclusive and 2 co-occurring mutated gene pairs. The already known ME gene pair of TP53-DROSHA is detected in the much smaller collection of the DKFZ Wilms tumors. Over half of the identified gene pairs represent new discoveries that have not been previously described in the literature. Four of them could be promising candidates for synthetic lethal genetic interactions.

**Conclusion:** These findings highlight the benefits of genetic interactions inference by exploring a different aspect of pediatric cancer data through pathways and propose new gene pairs for follow-up synthetic lethality experimentation.

## Background

Genetic interactions (GIs), the combination of two mutated genes that produces an unexpected phenotype, have been extensively investigated in cancer. In its most extreme form, these interactions can be either synthetic lethal (SL) or cooperative. Synthetic lethality is particularly interesting for uncovering new therapeutic targets by identifying cancer cell vulnerabilities. On the other hand, cooperation suggests an advantage for tumor cells and can offer insights into the mechanisms of cancer development.

The increased availability of sequenced cancer genomes has enabled the exploration of the genomic landscapes of tumors across many cancer types, giving the opportunity to perform in-silico studies to infer GIs [10]. Several inference methods leverage tumor mutations to test gene co-occurrence (CO) and mutual exclusivity (ME), testing the hypothesis whether two mutated genes occur together more or less often than expected [11–13,13,14]. Significantly co-occurring gene pairs may indicate cooperative GIs. Conversely, significantly mutually exclusive gene pairs may indicate synthetic sick or SL GIs. Key confounders in ME analysis include cancer subtypes and pathway epistasis, which often can result in findings that are not true synthetic sick/lethal candidates. Pathway epistasis, also known as functional redundancy, describes functionally related genes belonging to the same pathway, and mutating either of them leads to the same downstream effect. A mutation in any one of the genes is sufficient to alter pathway function, eliminating selective pressure for additional mutations in other genes within the same pathway [10,15,16]. As a consequence, the gene mutations will appear mutually exclusive in the CO/ME test but are likely not SL. While CO/ME testing identifies candidate gene pairs for SL and cooperation, it provides only statistical predictions and does not confirm functional relationships. Validation through literature review or experimental evidence is necessary to distinguish the biologically meaningful interactions from spurious associations and strengthen the confidence of the findings.

The application of GI inference methodologies in pediatric cancer continues to face challenges, as power limitations persist despite the expansion of collected cancer genomes, largely due to small sample sizes per cancer type [17,18]. However, pediatric cancer genomes remain a valuable resource for uncovering new biological insights. Incorporating prior knowledge can be instrumental for CO/ME testing. Biological pathways are summarized biological knowledge that aid in grouping functionally related genes. Aggregating genes in biologically meaningful functional groups or pathways, decreases the dimensionality of the data and contributes to increasing statistical power [19]. This can be a compelling way to perform an informed and biologically meaningful CO/ME testing to explore different aspects of the data.

Here, we present the pathway-informed genetic interactions framework (PIGI) applied to two pediatric cancer datasets, TARGET and DKFZ [20,21]. PIGI uses pathway knowledge from Reactome [22] to perform a biologically informed test, assesses CO and ME from pathway relationships and results in a set of co-occurring and mutually exclusive gene pairs. It uncovers new gene pair relationships and spotlights potential SL candidates by decreasing pathway epistasis, subtype effects, and cooperative interactions. Finally, PIGI reveals several ME gene pairs with an unknown relationship and four ME gene pairs, reflecting candidate SL GIs.

## Methods

### Data acquisition

WGS and WES data of TARGET (1669 primary tumors of 6 cancer types)[21] and DKFZ (961 primary tumors of 23 cancer types) [20] datasets were retrieved (mutation data are available to inspect in the following portals, https://www.cancer.gov/ccg/ and https://hgserver1.amc.nl/cgi-bin/r2/main.cgi?dscope=DKFZ_PED&option=about_dscope) and were processed with the same criteria as described in Daub et al. [17] to derive somatic SNVs and small indels with likely functional effects per tumor. The same criteria were applied, also, in excluding hypermutators; a hypermutator sample is considered to have more than 225 coding mutations.

### Reactome pathway database and pathway selection

Reactome is a curated and experimentally validated biological pathway database and, therefore, was selected as a pathway resource. The complete human pathway gene sets of version v86 and the corresponding pathway hierarchy was retrieved at https://reactome.org/download-data and all genes were used by their ENSEMBL IDs. Reactome groups its pathways in a hierarchical tree structure. The tree begins with level 0 (root) pathways that represent biologically broad and general functions. Moving deeper into the tree’s structure, level 1, 2, 3, …, x, the pathways become increasingly smaller and reflect greater biological specificity. Out of the 2647 pathways, a subset of pathways was selected to maintain statistical power, by excluding small pathways (< 10 genes) and large general pathways (> 300 genes) while still retaining biological relevance. The following criteria were applied: pathways at the second level in the tree that contain <300 and >10 genes were selected. Pathways that had fewer than 10 genes were omitted, whereas pathways that contained more than 300 genes were replaced by pathways of the next level deeper in the tree that satisfied the same rule (<300 & > 10 genes). This process was repeated iteratively until for each pathway the rule was satisfied. This resulted in a set of 413 pathways that contain 9460 genes.

#### Assignment of mutated genes to pathways

Each tumor’s mutated genes were assigned to their corresponding pathways. If a gene is mutated, then it is assumed that its corresponding pathway is also mutated. If a gene belongs to more than one pathway, it is assumed that all these pathways are mutated. This assignment resulted in a list of pathways that are mutated per tumor and per cancer type. To ensure higher confidence findings, each mutated pathway was retained in the test if it was mutated in 6 or more tumors. The final number of tumors tested via pathways per cancer type is shown in Suppl. Table 1.

### Co-occurrence & mutual exclusivity test

#### Pathway test with rediscover

Rediscover [23] was applied on mutated pathways per cancer type. Rediscover uses a statistical test that employs the Poisson-Binomial distribution to identify co-occurring and mutually exclusive events. Rediscover is a reimplementation of DISCOVER [11] and therefore, it takes into account the overall alteration rates of each tumor. This test was designed to detect genetic interactions on gene mutations but here we apply it to pathway mutations. Version 0.3.2 of “Rediscover” r package was used.

#### FDR estimation

Rediscover (in contrast to DISCOVER) does not perform an FDR control or estimation. FDR control methods were not suitable for this study due to the lack of continuous p-values and uniformity (p-value distribution skewed to 1). Therefore, an empirical FDR was estimated as described by Daub et al. 2021 [17]. This included calculating a null distribution by permuting the input matrices (nperm = 300), applying rediscover, and after obtaining the null distribution ng the found p-values were compared to the null distribution. The binary matrices were permuted using permatswap() function (mtype = “prab”, times = 300, rest default settings) of “vegan” r package version 2.6.4.

### Post-filtering

#### Removal of pathway pairs with shared genes

Shared genes between pathways led to identification of false significantly co-occurring pathway pairs, which would not have occurred without the shared genes. Therefore, all pathway pairs with at least one shared gene were removed from further analysis.

#### Omitting pathway pairs caused by mutation load effects

Mutation load differences among the tumors can yield false positives in mutual exclusivity testing. To account for these potential false positives, the Wilcoxon-Mann-Whitney test (wilcox.exact() function of r package “exactRankTests” version 0.8.35) was applied to the pathway mutation load of the tumors. Pathway mutation load was defined as the number of mutated genes (with functional impact on encoded protein) present in a given tumor that has a specific mutated pathway. If the mutation load of pathway A and pathway B were found significantly different (p-value <= 0.05) then they were excluded from the results.

#### Gene pairs retracing

For each pathway relationship found, the same relationship was applied to their underlying gene pairs. For example, pathway relationship A(genes: a, b, c)-B(genes: e, f) is mutually exclusive, therefore their corresponding genes are also marked as mutually exclusive: a-e, a-f, b-e, b-f, c-e, c-f. Per cancer type for each pathway pair these gene relationships were extracted (by omitting duplicates) to construct a network in Cytoscape.

#### Gene pair filtering

##### Filtering out gene pairs occurring in the same pathway

Gene pairs within the same pathway were excluded from the analysis. Gene pairs were assessed within the pathway subset that is used in this study.

##### Omitting mutually exclusive gene pairs caused by mutation load effects

Mutation load can confound the identification of mutually exclusive gene pairs, potentially resulting in false positives, as described above about pathway mutation load effect. To account for these potential false positives, for each gene pair, mutation load differences were assessed between tumors harboring mutations in each gene of the pair using a Wilcoxon-Mann-Whitney test. The pairs that have a p-value < 0.05 are omitted from the results.

##### Pathway-tumor mutation load threshold and gene pair filtering

For each cancer type, the distribution of tumor mutation load within each mutated pathway was calculated as the mutation load values vs the frequency of tumors with those values. The elbow point was identified from each distribution, and the mean of all elbow points across all pathways within a cancer type was calculated to establish a cancer-specific mutation frequency threshold. A global minimum threshold for gene mutation frequency was set to > =3. For cancer types where the mean elbow point for mutation frequency was below 3, gene pairs in which both genes have a mutation frequency > = 3 were retained. For cancer types where the cancer-specific mutation frequency threshold exceeded 3, this higher value was used as the threshold: only gene pairs with both genes meeting or exceeding that higher threshold were selected.

##### Gene pair selection for TARGET - B-ALL

For facilitating annotation, a subset of significant B-ALL gene pairs was selected due to the large number of findings. The subset consists of maximum 3 gene pairs per gene of the top 15 high mutation frequency genes found in the B-ALL tumors.

### Gene pair annotation strategy

Each gene pair was annotated using google scholar to find matching articles describing mutual exclusivity, co-occurrence, co-mutations, outcome, function of each gene, co-operation and synthetic lethality in the relevant cancer type they were found in or other cancer types.

## Results

### A pathway-informed genetic interaction framework to uncover genetic interactions in pediatric cancer

We developed the pathway-informed genetic interaction framework (PIGI) to detect candidate genetic interactions by assessing CO and ME of mutated pathways in pediatric cancers. Testing for genetic interactions at the pathway level increases statistical power and exploits data in a biologically meaningful way [22]. This is of particular importance for pediatric cancer, which, due to its relatively low incidence, has low sample numbers per tumor type. Furthermore, it allows easier interpretation of affected biological pathways, as well as reduces the number of detected pathway epistatic relationships [10,15,16]. Shortly, the PIGI framework involves four major steps (Fig. 1; Methods). First, data are processed using single-nucleotide variants (SNVs) and indels, resulting in a gene mutation profile for each tumor sample (Fig. 1, step 1). Then, these variants are matched with their corresponding biological pathways obtained from Reactome [22] to derive pathway mutation profiles per tumor. Second, pathway mutation profiles are then tested for CO and ME (Fig. 1, step 2). Rediscover examines the pathway mutation profiles per cancer type for significant CO and ME pathway relationships [11,23]. The third step includes retracing genes associated with the CO and ME pathway pairs, resulting in gene pairs that are either CO or ME (Fig.1, step 3). The final step involves filtering to prioritize the most significant CO and ME gene pairs (Fig.1, step 4). We then seek evidence in literature to propose potential biological explanations of the findings. Here, we apply the PIGI framework to investigate candidate genetic interactions in two pediatric cancer genome sequencing datasets, TARGET (Ma et al. 2018) and DKFZ (Gröbner et al. 2018). The TARGET dataset contains 1699 tumors across six cancer type and focuses mostly on leukemias (Suppl. Fig. 1). The DKFZ dataset is focused on central nervous tumors and comprises 23 cancer types amounting to 961 tumors in total (Suppl. Fig.1). Taken together, these datasets provide a diverse panel of tumors that allows for investigating the strengths of the PIGI framework and its ability to detect candidate genetic interactions.

**Figure 1.**
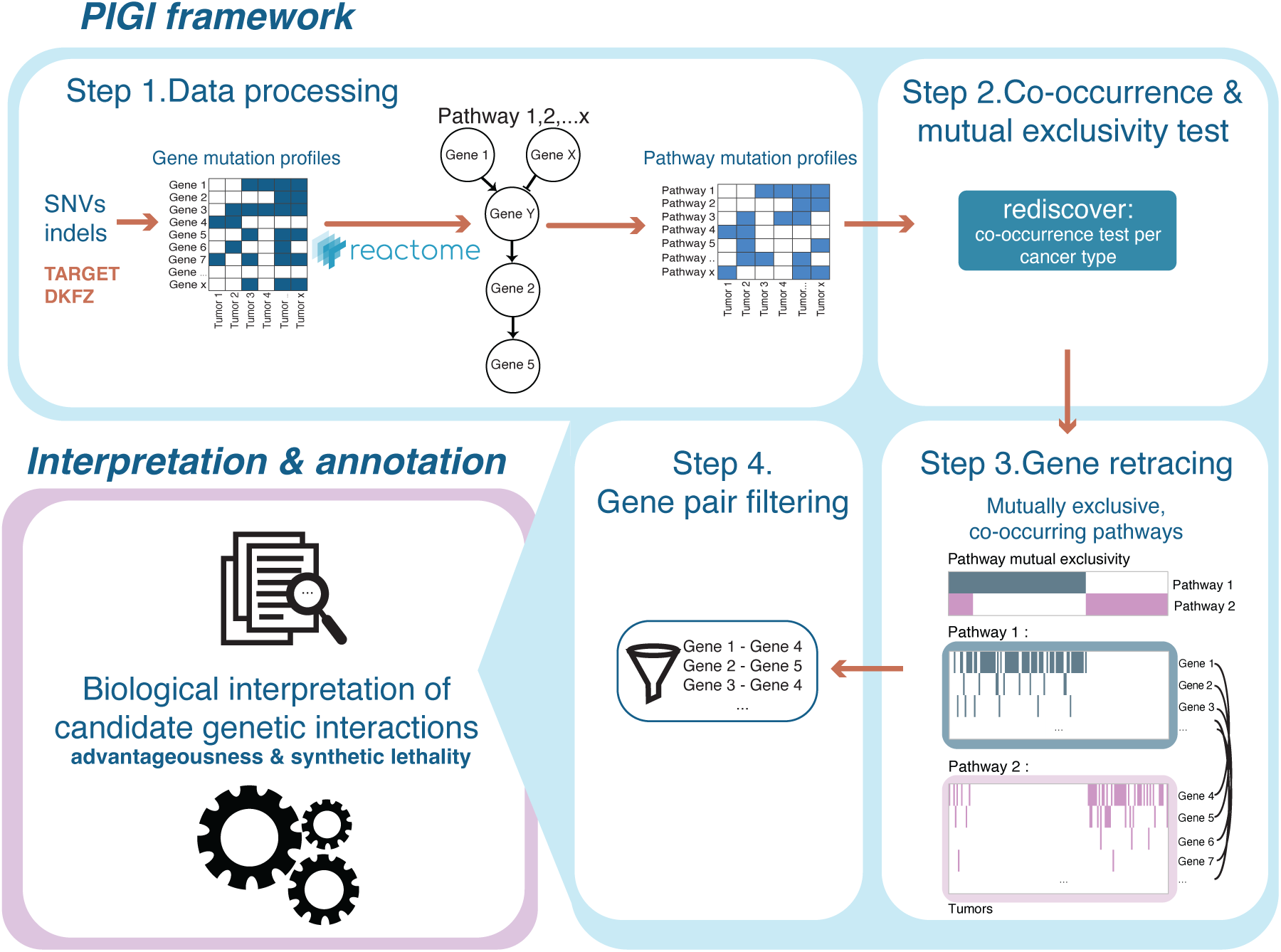
Overview of the PIGI framework. Schematic representation of the PIGI framework describing the steps for obtaining and interpreting ME and CO gene pairs. Step 1, processing of the data. Step 2, applying a co-occurrence test on pathway profiles. Step 3, associate genes with their corresponding pathways to extract ME and CO gene pairs. Step 4, filtering of gene pairs and finally, interpreting the resulting gene pairs using literature.

### Pathway ME and CO relationships are mostly cancer type specific

Applying the PIGI framework to the TARGET and DKFZ datasets revealed a high number of ME and several CO significant relationships between pathways (Fig. 2a, b, Suppl. Table 2a, b). In the TARGET dataset, PIGI detected 388 ME and 66 CO relationships between pathways (Fig. 2a). In DKFZ, 33 ME and 8 CO relationships are detected between pathways (Fig. 2b). As expected, most of the pathway pairs are directly or indirectly related to the hallmarks of cancer [24,25] such as proliferation-related signaling, DNA repair and cell death regulation. Individual pathways and pathway CO/ME pairs show similar properties to what has been found before for individual genes and CO/ME gene pairs regarding their association with certain cancer types [26–31]. First, individual pathways are associated either with specific cancer types such as “Notch-HLH transcription pathway” (B-cell Acute Lymphoblastic Leukemia (B-ALL), Fig. 2a) and “MHC class II antigen presentation” (Retinoblastoma (RB), Fig. 2b), or found across several cancer types, such as “NCAM signaling for neurite out-growth” (Fig. 2a) or “PIP3 activates AKT signalling” (Fig. 2b) [32]. Though overall, most individual pathways appear cancer-type specific. Second, CO/ME pathway pairs are, also, mostly cancer-type specific. Significant CO/ME pathway pairs are unique for each tested cancer type except for a few pairs (e.g. “Cellular Senescence” – “Costimulation by CD28 family”, “Cellular Senescence” – “Interferon Signaling” in Medulloblastoma-SHH (MB-SHH) and RB, Fig. 2b). This aligns with previous results for individual gene pairs which have also been found to be mostly cancer-type specific [11,17]. Third, we detect many more ME pathway pairs than CO pathway pairs (Suppl. Fig. 2a, b), similar to what has been reported before for gene pairs found in the two datasets [17]. We observe both CO/ME pathway pairs across most cancer types while in a few cancer types we find exclusively CO or ME pathway pairs (Suppl. Fig. 2a, b). A high number of ME pathway pairs are found for B-ALL and T-cell Acute Lymphoblastic Leukemia (T-ALL) tumors in TARGET, which is likely due to the large sample size (Fig.2a, Suppl. Fig.1a, 2a). The largest number of pathway pairs in DKFZ is found in retinoblastoma (RB) and High-grade glioma (HGG) (Suppl. Fig. 2b). Interestingly, Neuroblastoma (NBL), Osteosarcoma (OS), Medulloblastoma (MB), and Pilocytic astrocytoma (Fig. 2b, Suppl. Fig. 2a, 2b) had only few pathway pairs, despite having a considerable number of samples. This may, at least in part, be due to the exclusion of variant types other than SNVs and indels, and these tumor types may be driven by other events. For instance, larger structural variations in OS and MYCN amplifications in NBL. Taken together, these results show that most ME and CO pathway pairs follow previously found patterns in gene pairs and are mostly cancer-type specific.

**Figure 2.**
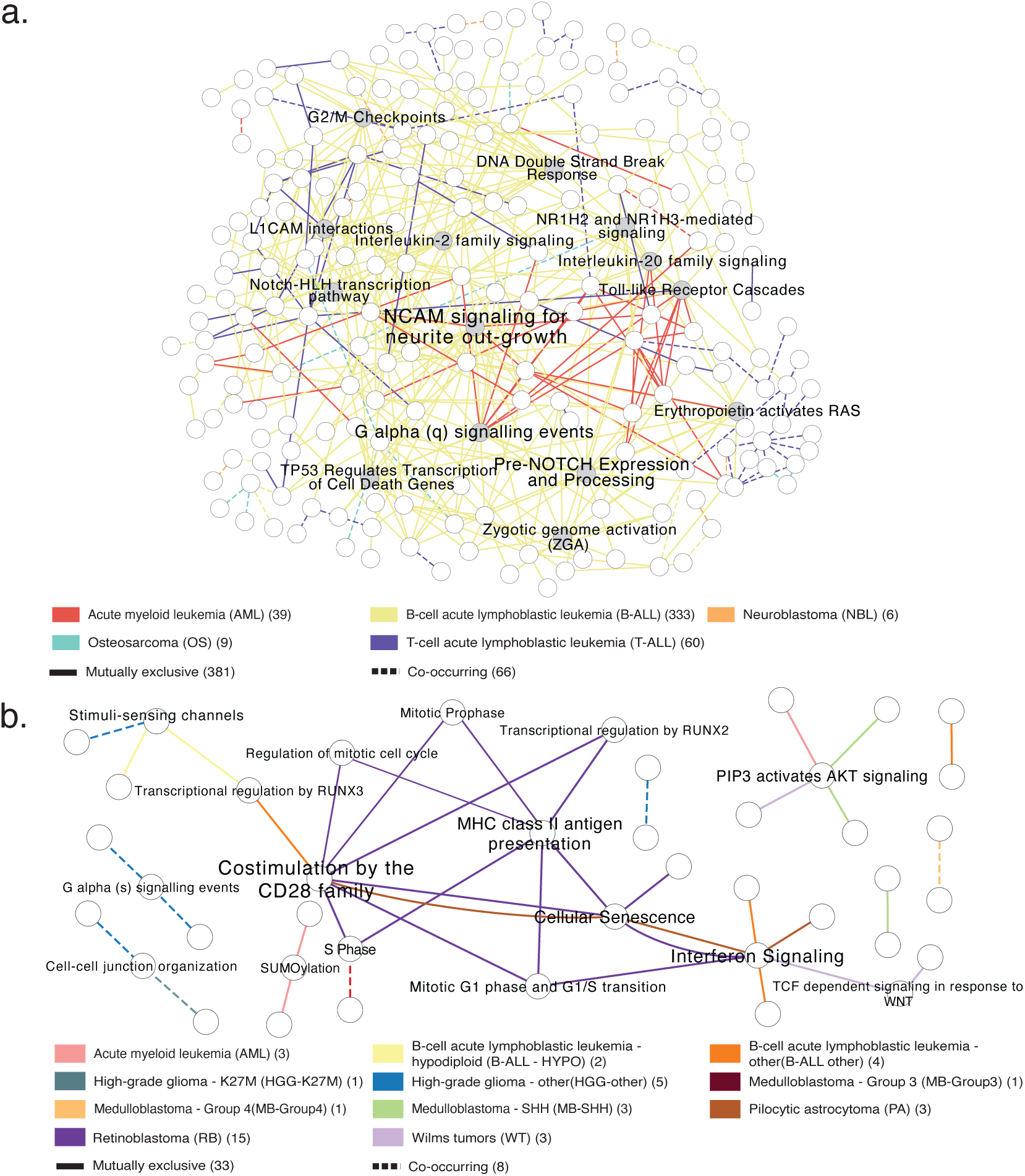
Significant pathway relationships detected by PIGI are largely cancer-type specific. Graph representation of mutually exclusive (ME) and co-occurring (CO) pathway pairs found by the PIGI framework in the TARGET and DKFZ datasets. Pathways are shown as nodes, ME/CO pathway pairs as solid/dashed lines respectively and colored by cancer type. The legend depicts the number of ME/CO pathway pairs in parentheses for each cancer type. a. ME & CO in TARGET tumors per cancer type, ME: 381 and CO: 66. Node labels (nodes in gray) shown only for pathways with 12 or more pathway pairs to prevent visual overload. b. ME & CO in DKFZ tumors, ME: 33 and CO: 8. Node labels shown only for pathways with two or more pathway pairs.

### PIGI controls for mutation load effects between pathways

The higher the mutation load of a tumor, the higher the chance that passenger mutations have accumulated in genes and their associated pathways. High mutation load differences between tumors can therefore lead to false detection of mutual exclusivity between genes and should be accounted for [33]. Similarly, mutation load differences between pathways likely exist. PIGI accounts for these by applying a Wilcoxon-Mann-Whitney test on the pathway mutation load between the tumors for which a particular pathway pair was found to be ME (Methods). Then, it retains only those pathway pairs without a significant difference in their mutation loads. By comparing the number of pathway pairs per cancer type with the corresponding cancer type-mutation loads (Suppl. Fig. 2a, c (TARGET) and 2b, d (DKFZ)), the number of pathway pairs per cancer type appears irrespective of the median tumor mutation load. B-ALL has a lower median mutation load compared to T-ALL, OS, and NBL, even though it has the highest number of ME gene pairs (Suppl. Fig. 2a, c). OS tumors show one of the highest median mutation load but have one of the lowest numbers of pathway pairs. Similarly, in DKFZ, HGG-K27M tumors show the highest median mutation load but have low number of pathways (Suppl. Fig. 2b, d). The highest number of pathway pairs is observed in RB which has one of lowest median mutation load. Overall, PIGI test sufficiently accounts for mutation load effects on pathway pairs.

### PIGI identifies several pathway-associated candidate genetic interactions

We next examined the results from the gene retracing step to identify individual CO and ME gene pairs (Fig. 3a, Suppl. Table 3). The gene pairs were retraced from CO/ME pathway pairs (FDR < 0.05), followed by removing gene pairs found in the same pathway and by applying a gene-mutation frequency threshold (see Methods section). We observe a considerably higher number of B-ALL gene pairs compared to the other cancer types, which are also exclusively ME (Fig. 3b). This likely reflects both the higher statistical power in B-ALL and the stringency of the p-value threshold. A more lenient threshold would benefit the detection of gene pairs in smaller cancer types (FDR < 0.2; Suppl. Table 4), albeit at the cost of increasing the number of likely false positives. Also, we observe a bias towards ME pairs, which has been observed before and can be attributed to primary tumors not being subjected to selective pressure, e.g., treatment [11]. It is also likely that the low number of co-occurrences is due to the aggregation of protein complex genes or functionally related genes that co-occur in pathways that provide an advantage to the cancer cells. This appears somewhat counterintuitive, as a single gene mutation is anticipated to be enough to disrupt a function; thus, gene mutations in these protein complexes or pathways are expected to appear mutually exclusive. That is not always the case, as we observe co-occurring gene pairs found previously by the CO/ME gene test (CO/ME-GT) on the same dataset by Daub et al. [17] (from now on referred as CO/ME-GT referring to the gene test results from this study) to be aggregated within pathways. For example, the co-occurring *JAK1-JAK3* in T-ALL tumors are aggregated within the “MAPK1/MAPK3 signaling” pathway (Suppl. Fig. 3a) and the *ATRX-H3F3A* in HGG-other tumors are aggregated within the “Telomere Maintenance” pathway (Suppl. Fig. 3b).

**Figure 3.**
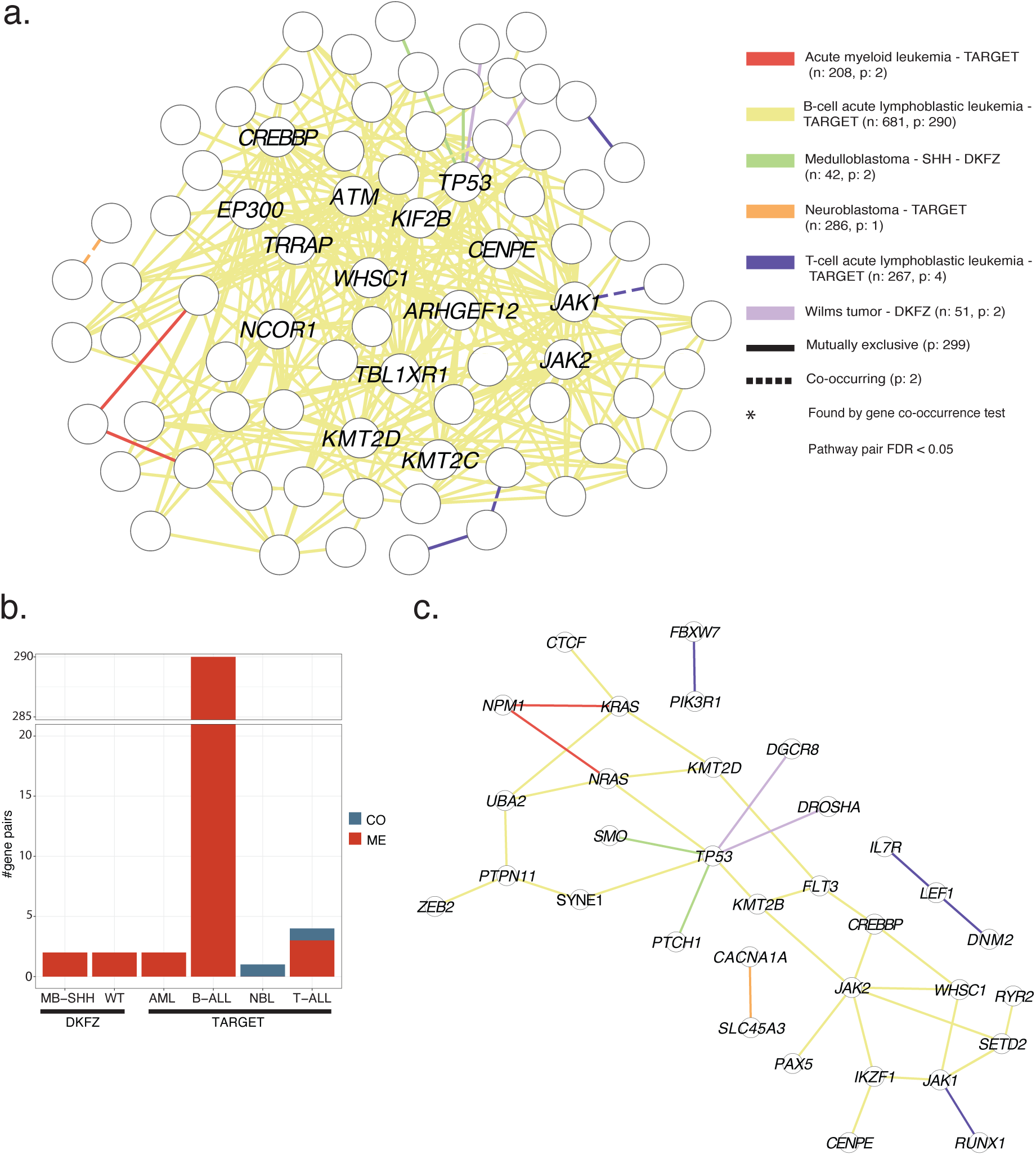
Co-occurring (CO) and mutually exclusive (ME) genes found by PIGI test in TARGET and DKFZ. a. Graph depicting CO and ME gene pairs found in tumors of both TARGET and DKFZ. Genes are shown as nodes, ME/CO gene pairs as solid/dashed lines respectively and colored by cancer type. The legend depicts the color per cancer type, shape for ME/CO, and in parentheses, the number of tested tumors and number of gene pairs per cancer type. b. Bar plot showing the numbers of CO and ME gene pairs found per cancer type in TARGET and DKFZ, c. Graph depicting CO and ME gene pairs (after prioritization of B-ALL findings) found in tumors of both TARGET and DKFZ. Legend as in a, except for the number of B-ALL gene pairs: 26.

Next, for annotation purposes, we restricted B-ALL gene pairs to the ones involving the top 15 highest mutation frequency genes (Methods). These results (Fig. 3c) consisted largely of new gene pairs and candidate genetic interactions that have not been observed before in these datasets in CO/ME-GT gene pairs [17]. The only previously detected pair is the *TP53-DROSHA* ME in Wilms tumors (WT), which was observed in 111 TARGET Wilms tumors. Whereas here, we detected *TP53-DROSHA* ME in DKFZ WTs, a much smaller tumor set of only 40 tumors. Therefore, this shows that PIGI was able to retrieve this finding with ∼60% fewer tumor samples. Interestingly, PIGI does not detect a significant *TP53-DROSHA* ME in TARGET WTs. This is likely due to the number of tumors displaying mutual exclusivity in TARGET and DKFZ, which may not be sufficient to reach statistical significance under the current FDR thresholds. To investigate this further, we compared the number of tumors supporting the *TP53-DROSHA* ME as detected by PIGI and CO/ME-GT in both TARGET and DKFZ (Suppl. Fig. 4a, b, c, d) [17]. In TARGET, CO/ME-GT found TP53-DROSHA ME supported by 31/115 tumors (∼27%) (Suppl. Fig. 4a), with complete mutual exclusivity and an FDR ranging from 0.0867 to 0.1033. In contrast, PIGI identified TP53-DROSHA related pathways (here presenting the ME pathways with the lowest FDR detected) in 44/110 tumors (∼40%), with two tumors overlapping and FDR of 0.143. This shows that, despite the increase in tumors supporting the mutual exclusivity, the increase is not large enough to reach statistical significance. On the other hand, CO/ME-GT in DKFZ tested *TP53-DROSHA* ME in 8/44 tumors (∼18%) and was not considered significant (FDR > 0.2). While PIGI detected *TP53-DROSHA* ME through pathways in 19/33 tumors (∼56%) with pathway FDR of 0.0031. Here, the substantial increase in ME supporting tumors – more than half of the dataset – was sufficient to reach statistical significance. Taken together, these findings highlight how pathway-level analysis with PIGI can amplify ME signals in small datasets such as DKFZ WT and underscore the importance of the chosen FDR threshold. A more lenient threshold, as used by CO/ME-GT, allows PIGI to detect significantly ME pathways that contain *TP53-DROSHA* in TARGET WT. However, these results are disregarded from the downstream analysis due to significantly different mutation load between the tumors of the two pathways. Albeit a sole case of recovering known findings in smaller datasets, it indicates the potential of using pathways for identifying genetic interactions in smaller datasets.

### A prioritized set of gene pairs reveals candidate synthetic lethal genetic interactions

The literature search revealed several candidate synthetic lethal gene pairs (Fig. 4 and Suppl. Table 5). We grouped CO/ME gene pairs according to their likely biological explanation: 4 candidate synthetic sick/lethal, 20 unknown, 7 pathway epistasis, 4 false positives, and 2 cancer subtype. The candidate synthetic sick/lethal genetic interactions consist of *TP53-DROSHA*, observed before in CO/ME-GT [17], the new *TP53-DGCR8* pair in WT, and *CREBBP-WHSC1* and *SYNE1-TP53* in B-ALL. The candidate synthetic lethal gene pairs represent ∼11% (4/37) of the found CO/ME gene pairs by PIGI, only slightly lower than the ∼13% (3/24) found in CO/ME-GT [17] for both datasets combined. The largest proportion of the gene pairs lacks supportive literature evidence. Thus, their biological explanation remains unknown and warrants further investigation. Frequently mutated genes such as *TP53*, *KRAS*, and *NRAS* are mutually exclusive with less often mutated genes such as *RYR1* and *UBA2*. This could be an artifact caused by mutation load differences, since long genes (gene length > 100kb) were included in the study. However, that is not the case for most of the findings. For the majority of gene pairs, the Wilcoxon-Mann-Whitney p-value for mutation load differences is above 0.1. Only 32/301 gene pairs have a p-value between 0.1 and 0.05 that could warrant further attention regarding their mutation loads. On the other hand, these low-frequency genes are relatively understudied within the context of the cancer type in which they were found. Consequently, almost no information is available about the relationship of these gene pairs leading to an inconclusive biological explanation. For example, *KRAS* and *CTCF* are mutually exclusive and have largely different biological roles according to literature. While *KRAS* is well-known for its role in MAPK and PI3K signaling and thereby regulating proliferation [34,35], *CTCF* is involved in regulating 3D genome organization, mRNA splicing and, DNA recombination [36]. These processes are not directly related to each other and therefore cannot explain their potential interaction. Few ME gene pairs (n=2) can likely be explained through cancer subtype (2/35, ∼6%), a number much lower than the ∼38% (9/24) proposed before [17]. This may be due to the aggregation of genes into pathways. For example, in B-ALL low-hypodiploid subtype, *TP53*, *RB1* and *IKZF2* are frequently mutated, whereas in the near-haploid B-ALL subtype, *IKZF3* and genes involved in RTK and RAS signaling are targeted. Despite both having different genetic alterations, both have been shown to display activation of the RAS and PI3K signaling pathways [37]. If tumors from these two subtypes are included in PIGI, due their shared pathway effects, they would be grouped in the pathway analysis, subsequently removing the potential ME confounded by their subtypes. When examining the results of PIGI and CO/ME-GT [17], *KRAS-TP53*, previously found to be ME in the CO/ME-GT [17] and each gene mutation being associated with different cancer subtype, is now aggregated through the pathway ‘Transcriptional regulation by RUNX3’ (Suppl. Fig. 5) and no longer detected as ME by PIGI. Thus, pathway-level aggregation in PIGI can resolve some of the false mutual exclusivity caused by subtype-specific gene mutations. Overall, PIGI indicated four new candidate synthetic lethal genetic interactions and 20 new gene pairs with, to our knowledge, roles yet to be discovered.

**Figure 4.**
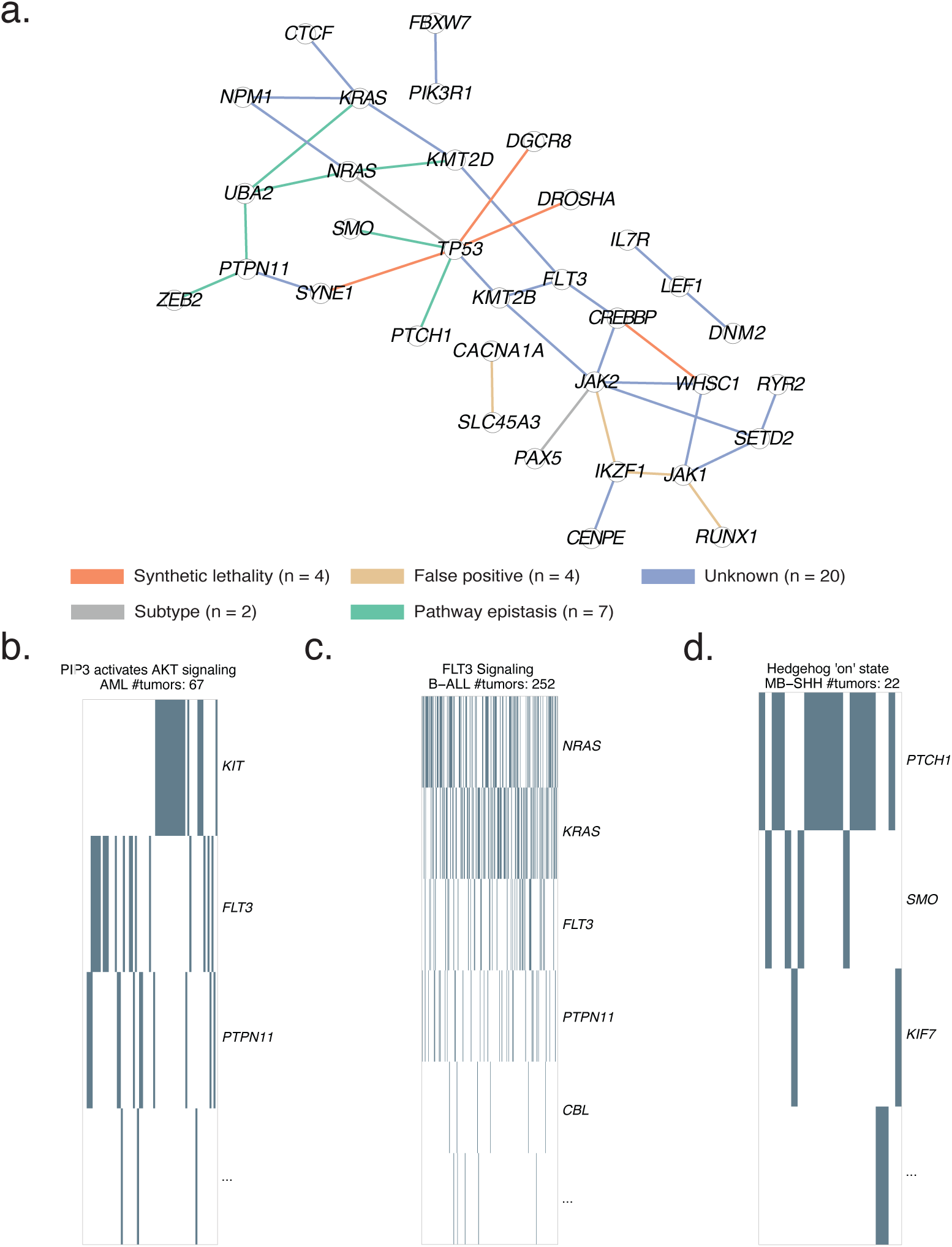
Biological interpretation of prioritized CO/ME gene pairs found in tumors of TARGET & DKFZ and examples of PIGI decreasing pathway epistasis. a. Graph representation of CO and ME gene pairs demonstrating their likely biological interpretation: synthetic lethality (red), false positive (brown), unknown (blue), subtype (grey), or pathway epistasis (green). b. Heatmap of “PIP3 activates AKT signaling” pathway in AML tumors (TARGET) showing mutual exclusivity of FLT3-KIT within the pathway. c. Heatmap of the FLT3 Signaling pathway in B-ALL tumors (TARGET) showing mutual exclusivity between FLT3-KRAS. d. Heatmap of the “Hedgehog ‘on’ state” in MB-SHH tumors (DKFZ) showing mutual exclusivity between PTCH1-SMO.

### Pathways in PIGI framework decrease pathway epistasis

Pathway epistasis, where the same downstream functional effect is caused by mutating individual genes within the same pathway, often confounds mutual exclusivity analyses when searching for synthetic lethality [10,15,38]. Despite PIGI employing pathways, ME findings (Fig. 4a) still contain a few gene pairs characterized by pathway epistasis, e.g., *TP53-PTCH1* in MB-SHH [39], *NRAS-KMT2D* [40], *PTPN11-UBA2* in B-ALL [41]. This reflects the incompleteness of the pathway database in recording all genes associated with a particular pathway, and pathway epistasis also assumes that pathways are linear with no crosstalk between pathways. Therefore, employing pathways will not completely avoid pathway epistasis. Still, we observe a decrease of pathway epistasis findings, ∼20% (7/35) compared to the ∼42% (10/24 ME gene pairs) pathway epistasis gene pairs observed in Daub et al. [17]. Three examples of pathway epistasis no longer confounding mutual exclusivity analyses are depicted in Fig. 4b, c, d. *FLT3-KIT* (Acute Myeloid Leukemia (AML), Fig. 4b), *FLT3-KRAS* (B-ALL, Fig. 4c), and *PTCH1-SMO* (MB-SSH, Fig. 4d) all show clear mutual exclusivity within the same pathway. PIGI removes these ME findings by aggregating individual genes within pathways, thereby decreasing the confounding effects of pathway epistasis when searching for synthetic sick/lethal gene pairs.

### New candidate synthetic lethal genetic interactions involving oncogenes/tumor suppressors genes or DNA repair genes

Investigating vulnerabilities, such as those arising from synthetic lethal genetic interactions within DNA damage repair (DDR) pathways, has enabled the development of new targeted therapies. This approach has led to the breakthrough discovery of PARP inhibitors, which exploit defects in homologous recombination, but also other inhibitors for *ATR*, *ATM*, *WEE1,* and *PRMT*[42–47]. Therefore, to identify likely candidates for synthetic lethal genetic interactions, we investigated PIGI findings (*TP53-DROSHA*, *TP53-DGCR8*, *SYNE1-TP53*, *CREBBP-WHSC1*) in the literature and their relation to DDR. These genes are implicated directly or indirectly in DDR, and it is probable that mutations in these gene pairs can lead to a synthetic sick/lethal phenotype. Also, some of these genes are known tumor suppressor genes (*TP53*, *CREBBP*) or oncogenes (*WHSC1*).

Here, we postulate a literature-based biological hypothesis of the function of these candidate synthetic sick/lethal genetic interactions. *TP53* and *DROSHA* play an important role in DDR. *TP53* controls G1 to S cell cycle progression, and loss of *TP53* is associated with a higher number of double-stranded breaks (DSBs) [48]. *DROSHA* plays, in addition to its well-described function in microRNA processing, a role in DDR pathways by processing DNA-damage-induced long non-coding RNAs [49–51]. Loss of either *DROSHA* in combination with loss of *TP53*, could increase the number of DSBs to a state that can be intolerable and fatal for the cell (Fig. 5a). The same effect is expected for *TP53-DGCR8*, as *DGCR8* is part of the microprocessor complex together with *DROSHA* [50,52], and loss of *DGCR8* could potentially disrupt the functionality of this protein complex. Secondly, *SYNE1* and *TP53* constitute another case of potential SL. *SYNE1* is part of the LINC complex which is important for correct DNA repair by surrounding the damaged DNA locus with a nuclear envelope fold (Fig. 5b) [53–55]. We anticipate that upon loss of *TP53* and *SYNE1,* DNA repair is hampered, possibly leading to synthetic sickness or lethality. Finally, *CREBBP* and *WHSC1* are both playing an important role in DDR. *WHSC1* is an oncogene that initiates non-homologous end joining and is involved in epithelial-to-mesenchymal transition [56]. Whereas *CREBBP* is a direct regulator of DDR [57]. Due to their substantial role in DDR, targeting both genes could result in high levels of unrepaired DNA and subsequently a synthetic sick/lethal effect (Suppl. Fig. 6). Taken together, these examples highlight the potential of detecting likely synthetic sick/lethal relationships with PIGI, although follow up experimentation is necessary to assess the validity of these hypotheses.

**Figure 5.**
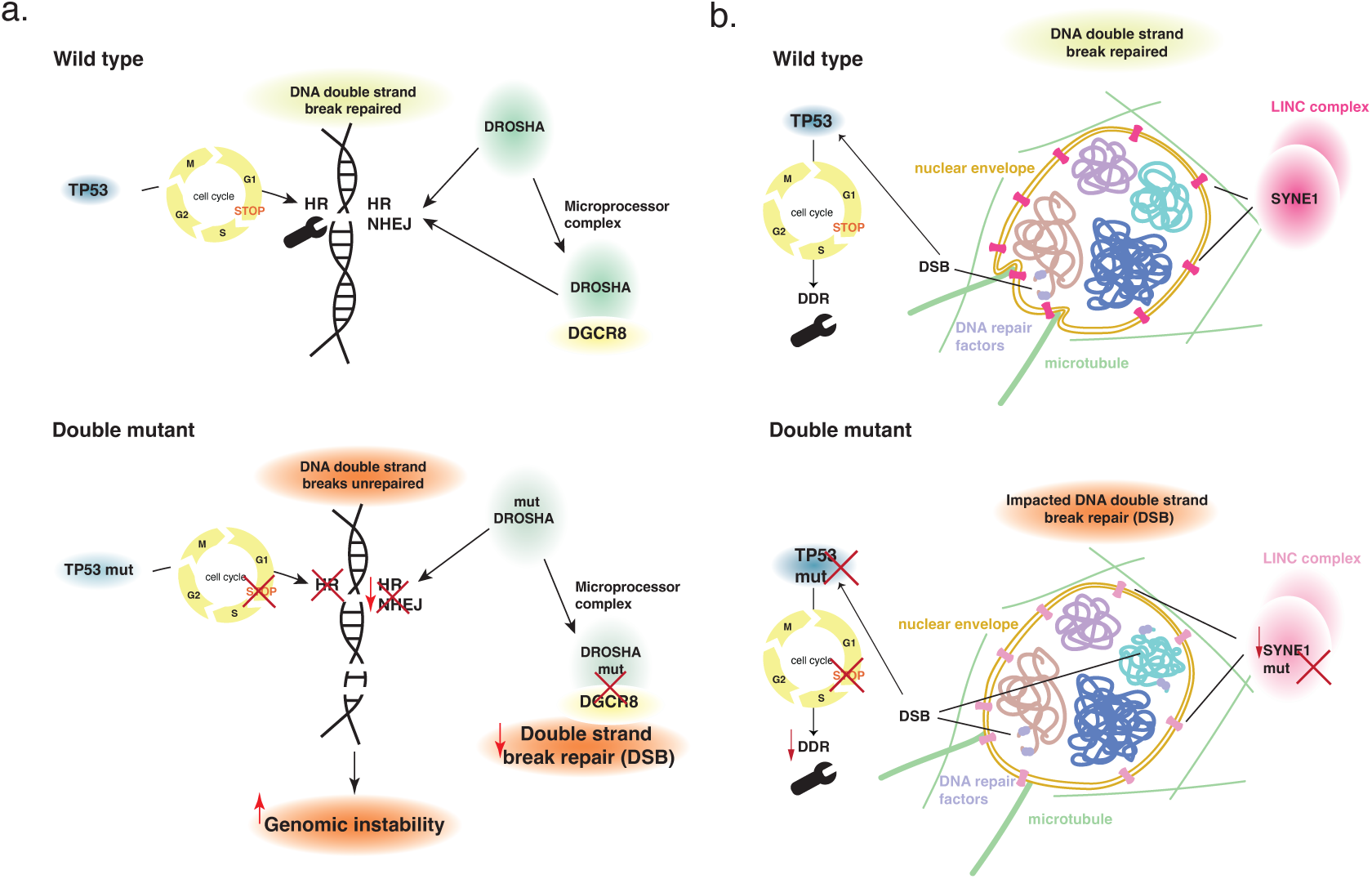
Biological interpretation of candidate synthetic lethal gene pairs found by PIGI test. a. Top: function of wild-type TP53, DROSHA in DNA strand break repair processes; homologous recombination (HR) and non-homologous end joining (NHEJ). Bottom: loss-of-function TP53, DROSHA and their effect in DNA damage repair processes could lead to increased accumulation of double-strand breaks and genomic instability. b. Top: wild-type function of TP53 and SYNE1 via the LINC complex in DNA damage repair (DDR). Bottom: loss-of-function TP53, SYNE1, and their effect in DDR processes. Failure to pause cell cycle progression in combination with suboptimal nuclear mechanics could lead to increased accumulation of unrepaired DNA. Part of the figure is adapted from Lottersberger et al. 2015.

Overall, PIGI framework successfully identified new mutually exclusive (ME) gene pairs in the two primary pediatric cancer datasets, TARGET and DKFZ. It uncovered new gene pair relationships, spotlighted potential SL candidates by decreasing pathway epistasis, subtype effects, and cooperative interactions. It detected a known ME gene pair in the much smaller dataset of the DKFZ Wilms tumors. Finally, PIGI revealed several ME gene pairs of unknown roles and four ME gene pairs that are promising candidate SL genetic interactions.

## Discussion

Here we present the pathway-informed genetic interaction framework (PIGI) to leverage tumor mutations in inferring candidate GIs. To our knowledge, this is the first study that employs mutated pathways to infer genetic interactions in pediatric cancer. The PIGI framework employs a statistical test to explore ME and CO between pairs of pathways to retrieve CO and ME relationships between gene pairs. The framework detected 35 ME and 2 CO relationships between gene pairs across 29 cancer types. PIGI spotlighted potential SL candidates by decreasing pathway epistasis, subtype effects, and cooperative interactions. Supporting evidence indicated 4 gene pairs as candidate synthetic lethal, *CREBBP-WHSC1*, *SYNE1-TP53* in B-ALL, and *TP53-DROSHA*, *TP53-DGCR8* in WT, and 20 gene pairs of unknown impact.

PIGI provides a first analysis that exploits pathways for genetic interaction inference in pediatric cancer. Several studies have previously used prior knowledge, such as pathways or protein-protein/functional networks, to detect genetic interactions in adult cancer [11,15,58,59]. Among these approaches, Mutex and BriDGE utilize pathways. Mutex employs signaling pathways to detect gene sets with a shared downstream effect and identify mutual exclusivity in TCGA cancer genomes [58]. BriDGE incorporates a large selection of pathways (833) from KEGG, Biocarta and Reactome to identify genetic interactions in genome-wide association studies [59]. PIGI, however, uses a set of 413 pathways that cover many biological processes to detect candidate genetic interactions. Also, this set of pathways is comprised only by Reactome to ensure up-to-date pathways are included which are recorded in the same consistent and systematic manner. While the aforementioned studies focused on adult cancer, PIGI was designed to investigate pediatric cancer genomes which often face limitations in statistical power. The shorter but comprehensive set of pathways in the PIGI framework aids in decreasing the number of tested variables while still decreasing pathway epistasis and cancer subtype effects. Essentially, PIGI provides a biologically informed testing approach towards ME to identify potentially synthetic sick/lethal genetic interactions.

Pathways confer advantages in pathway analyses, but they also entail limitations. Results are dependent on the completeness of the pathways and their definition. Crosstalk or tissue-specific functional relationships are not always adequately recorded [10,22,60]. Consequently, this still retains some pathway epistasis ME with the PIGI framework. Another challenge is the lack of dedicated pathway resources describing the processes across different developmental stages and ages to describe pediatric cancer, a disease stemming from developing tissues [61,62]. One way to cope with the aforementioned challenges is to retain only active pathways within each tumor by inspecting transcriptome data. However, this analysis can be challenging due to the uncertainty about to what extent developmental pathways in Reactome are representative of human developing tissues. Finally, crosstalk between pathways leads to several pathways having overlapping genes, e.g., TP53 is present in several pathways in Reactome. Albeit biologically sound, it can lead to statistical inflation and incorrect findings. The PIGI framework accounts for that by applying strict thresholds, removing ME/CO pathways with overlapping genes and ME genes that are part of the same pathway.

Confounders pose an extra challenge in the PIGI framework and other similar methods. Cancer subtypes confound mutual exclusivity testing, leading to false candidate synthetic lethal pairs. This can be solved by performing a mutual exclusivity test per cancer subtype [10]. However, this solution is not well-suited for pediatric cancer datasets due to the already limited number of samples per cancer type. The PIGI framework decreases cancer subtype effects by grouping the subtype-associated mutated genes into their common pathway. Conversely, tumor mutation load can affect mutual exclusivity detection. Applying a mutual exclusivity test to tumors with high differences between their mutation loads leads to false-positive mutually exclusive gene findings [33]. Likewise, mutated pathways in tumors with high mutation load will falsely appear as significantly mutually exclusive with pathways derived from tumors with lower mutation load. Unlike BriDGE and Mutex, PIGI controls mutation load effects for both pathway and gene pairs by applying a Wilcoxon-Mann-Whitney test between the tumors of the two pathways or genes.

Completeness of pediatric cancer datasets is a significant parameter in applying genetic interaction inference methodologies. A low level of dataset completeness leads to low sample sizes. Thus, combining pediatric cancer datasets can be beneficial in increasing dataset completeness and, consequently, uncovering new candidate genetic interactions in rare cancer types. However, combining datasets can introduce confounders, such as variations in experimental protocols and analysis pipelines, making it crucial to account for these differences during analysis. Exploring other layers of the mutational landscape of pediatric cancer may also be informative to increase dataset completeness. SNVs and indels constitute an incomplete representation of the mutational landscape within tumor cells, especially for cancers mainly driven by copy number variants (CNVs) or structural variants (SVs). Gradually, more pediatric cancer studies have been integrating SVs and CNVs in ME testing [63,64] as it confers an advantage of describing the complete mutational landscape of each cancer type and increasing the chance of revealing candidate SL genetic interactions in more cancer types. We deem that gene variant integration will be valuable, not only in applying PIGI or other pathway CO/ME testing methods but also on gene-level CO/ME testing, in maximizing the discovery potential of candidate SL genetic interactions in pediatric cancer.

## Conclusions

In summary, PIGI is a valuable testing framework for studying pediatric cancer genomes to identify new candidate synthetic lethal genetic interactions using publicly available data. While genome-wide screens of tumor-derived cell lines are the only direct way to detect genetic interactions, PIGI provides candidate tumor-derived genetic interactions for guided, follow-up experimentation in pediatric cancer.

## Abbreviations

AML: Acute Myeloid leukemia
B-ALL: B-cell Acute Lymphoblastic Leukemia
CO: co-occurrence
CNV: Copy Number Variant
DDR: DNA Damage Repair
DSB: Double Strand Break
FDR: False Discovery Rate
GI: Genetic Interaction
HGG-other: High-grade glioma – other
MB: Medulloblastoma
MB-SHH: Medulloblastoma – SHH
ME: mutual exclusivity
NBL: Neuroblastoma
OS: Osteosarcoma
PIGI: Pathway-Informed Genetic Interaction framework
RB: Retinoblastoma
SL: Synthetic Lethal
SNV: Single Nucleotide Variant
SV: Structural Variant
T-ALL: T-cell Acute Lymphoblastic Leukemia
WGS: Whole Genome Sequencing
WES: Whole Exome Sequencing
WT: Wilms tumor

## Declarations

### Ethics approval and consent to participate

The Data Access Committees of TARGET and DKFZ datasets have approved the sharing and use of the tumor data for this study. Accession codes for DKFZ: RP012816, PRJEB11430 (European Nucleotide Archive); EGAS00001001139, EGAS00001001953, EGAS00001000607, EGAS00001000381, EGAS00001000906, EGAS00001001297, EGAS00001000443, EGAS00001000213, EGAS00001000263, EGAS00001000192, EGAS00001000255, EGAS00001000254, EGAS00001000253, EGAS00001000256, EGAS00001000246, EGAS00001000379, EGAS00001000380, EGAS00001000346, EGAS00001000349, EGAS00001000347, EGAS00001000192 (European Genome-Phenome Archive). Study identifiers for TARGET: phs000218, phs000463, phs000464, phs000465, phs000467, phs000471, and phs000468.

### Consent for publication

Not applicable.

### Availability of data and materials

All data are available with accession code/ identifiers as indicated in Ethics approval section. All mutation data of TARGET and DKFZ datasets can be inspected in their respective portals https://www.cancer.gov/ccg/ and https://hgserver1.amc.nl/cgi-bin/r2/main.cgi?dscope=DKFZ_PED&option=about_dscope. Code availability on: https://doi.org/10.5281/zenodo.15881284

### Authors’ contributions

Conceptualization: A.S., J.D., P.K., Methodology: A.S., R.G., P.K., Analysis: A.S., Supervision: J.D., P.K. Manuscript writing: A.S., P.K., Manuscript revision: A.S., R.G., J.D., P.K. All authors have read and approved the final version.

## Acknowledgements

The authors thank Puck Veen, Femke Albertsboer, Joanna von Berg and fellow lab members for insightful discussions and helpful feedback.

## Authors’ information (optional)

Princess Máxima Center for Pediatric Oncology, Utrecht, The Netherlands: A.S., R.G., J.D., P.K.

Oncode Institute, Utrecht, The Netherlands: J.D.

Center for Molecular Medicine, UMC Utrecht and Utrecht University, Utrecht, The Netherlands: P.K.

## Corresponding author

Correspondence to

