## Supplementary Figures for "A pathway-informed mutual exclusivity framework to detect genetic interactions in pediatric cancer"

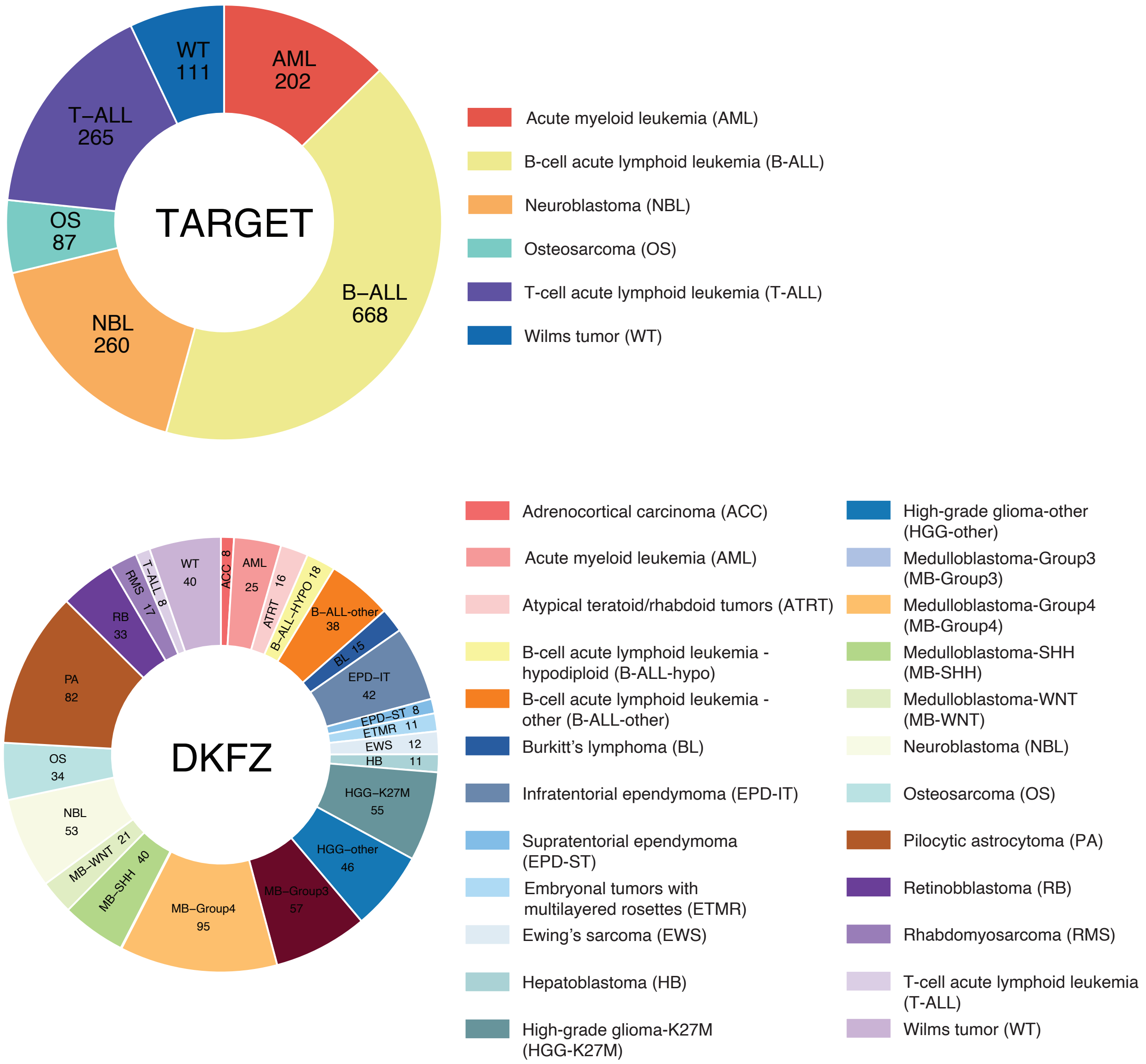


**Supplementary figure 1. Overview of number of tumors per cancer type in TARGET (up) and DKFZ (down) that were used in this study after sample filtering (omitting relapse, hypermutator tumors).**

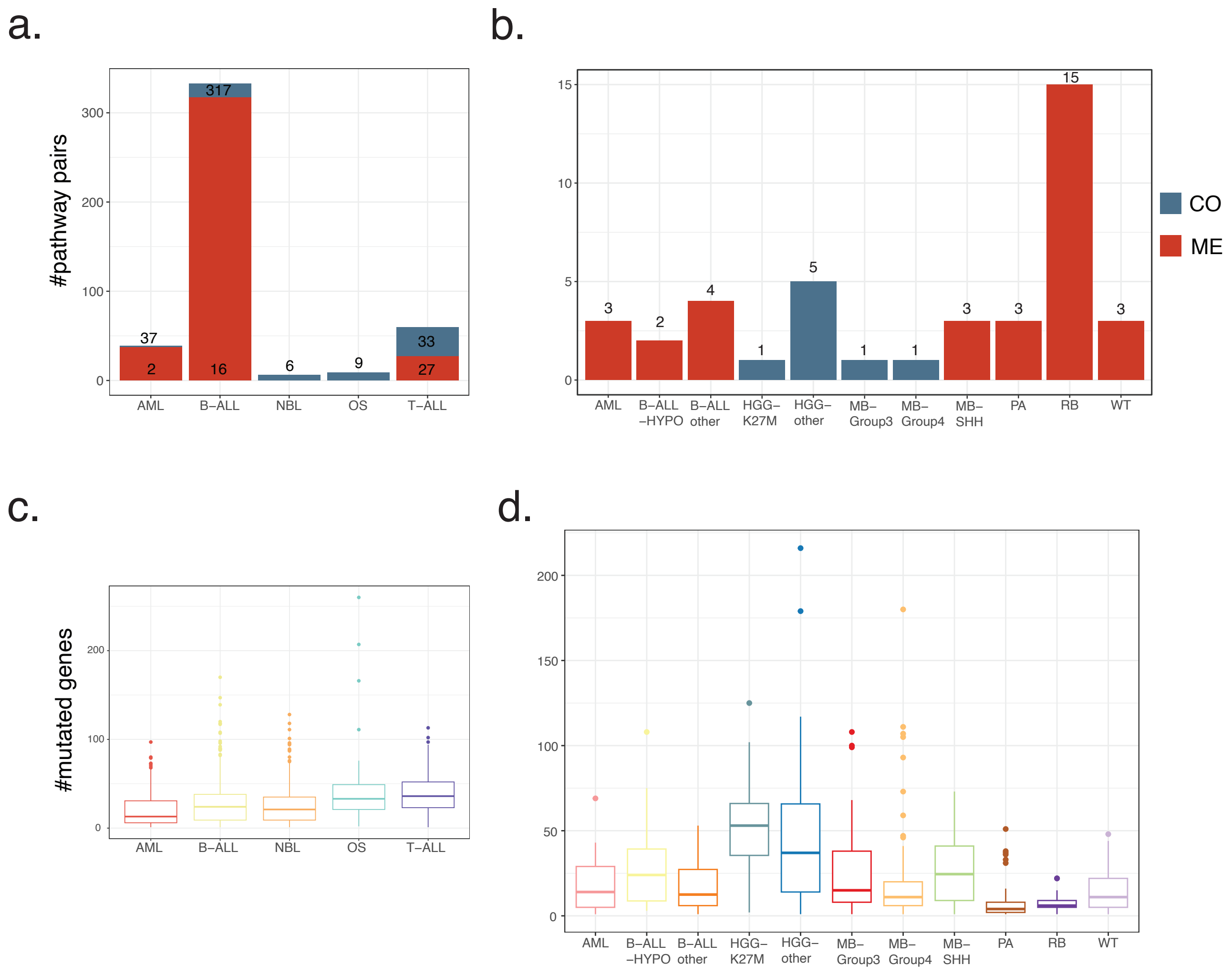


**Supplementary figure 2. Number of ME/CO pathways and median mutation count per cancer type.** a. number of ME & CO pathway relationships in TARGET tumors, ME: 381 and CO: 66, b. number of ME & CO pathway relationships in DKFZ tumors, ME: 33 and CO: 8, c. boxplot of number of mutated genes per tumor per cancer type in TARGET tumors, d. boxplot of number of mutated genes per tumor per cancer type in DKFZ tumors.

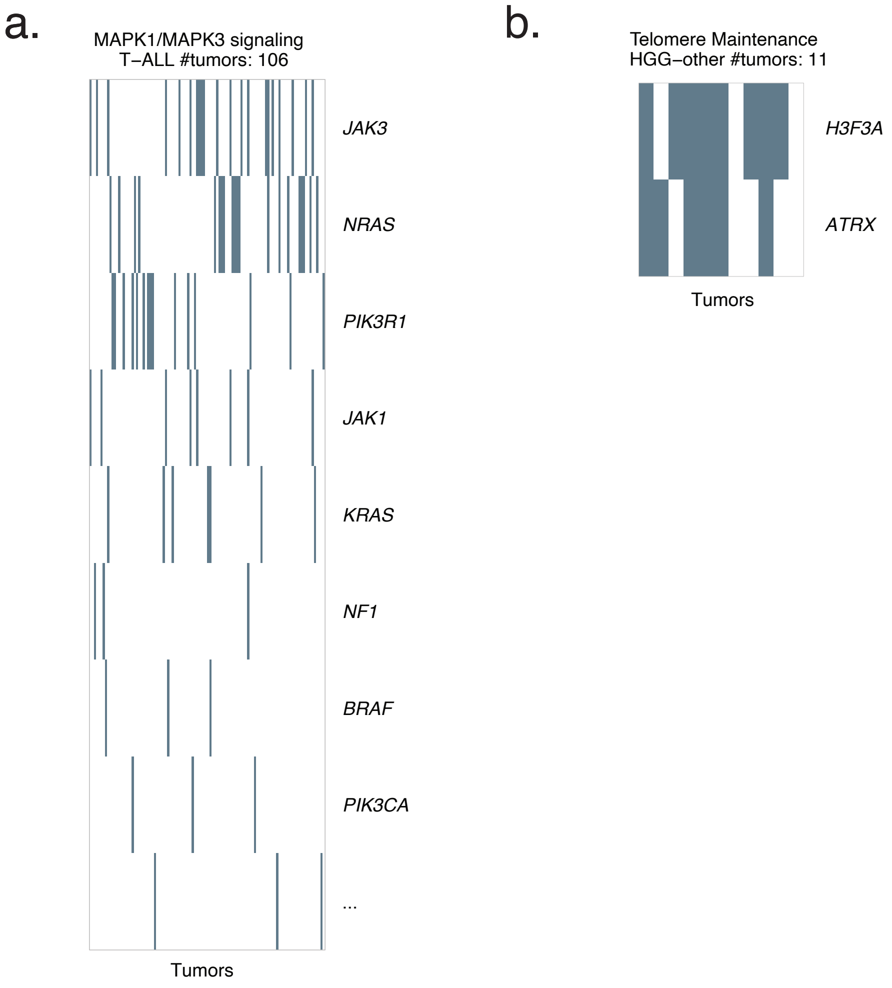


**Supplementary figure 3. Pathway-gene mutation heatmaps showing within pathway gene co-occurrence.** a. Heatmap of mutated genes in MAPK1/MAPK3 signaling pathway in TARGET T-ALL tumors (#tumors: 106) showing the co-occurrence of *JAK1* & *JAK3*. b. Heatmap of mutated genes in Telomere Maintenance pathway in DKFZ HGG-other (#tumors: 11) showing the co-occurrence of *H3F3A* & *ATRX*.

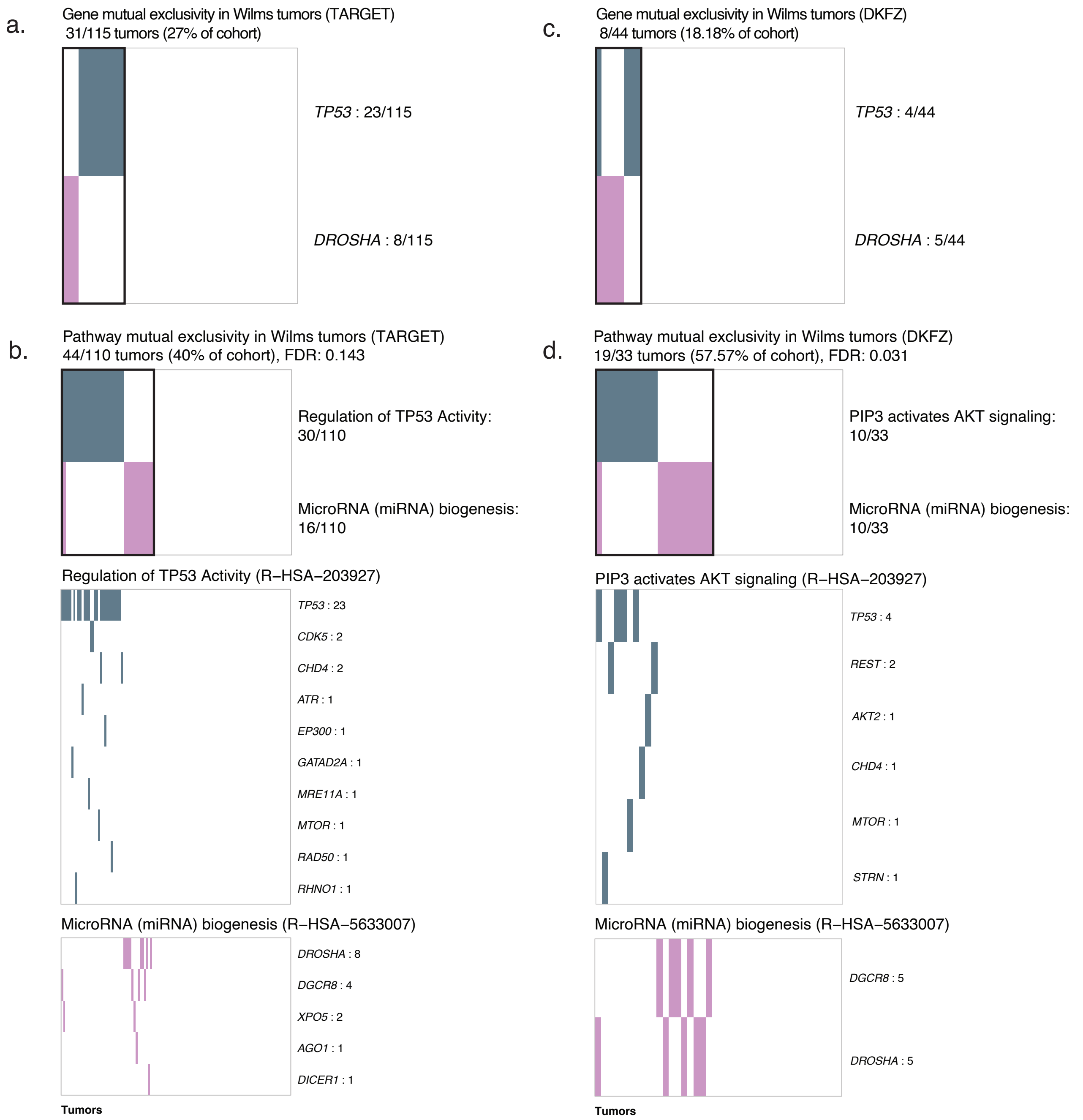


**Supplementary figure 4. Gene and pathway-gene mutation heatmaps presenting mutual exclusivity between TP53 and DROSHA across DKFZ and TARGET Wilms tumors.** a. Gene mutation heatmap of *TP53* and *DROSHA* in TARGET Wilms tumor (WT). Each row describes with color the gene that is mutated in the tumors and each label contains the number of mutated tumors out of the whole Wilms tumor dataset. b. Pathway-gene mutation heatmap showing pathway mutual exclusivity between TARGET Wilms tumors with mutated 'Regulation of TP53 Activity' (*TP53* related) and 'MicroRNA (miRNA) biogenesis' (*DROSHA* related) pathways and the underlying mutated genes for each pathway per tumor in the cohort. Each row describes with color the pathway/ gene that is mutated in the tumors and each label contains the number of mutated tumors out of the whole Wilms tumor dataset. c. Gene mutation heatmap of *TP53* and *DROSHA* in DKFZ Wilms tumor (WT). Rows describe the same as in a. d. Pathway-gene mutation heatmap showing pathway mutual exclusivity between DKFZ Wilms tumors with mutated 'PIP3 activates AKT signaling' (*TP53* related) and 'MicroRNA (miRNA) biogenesis' (*DROSHA* related) pathways and the underlying mutated genes for each pathway per tumor in the cohort. Rows describe the same as in b.

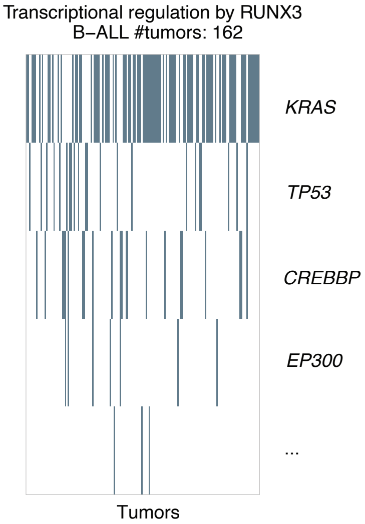


**Supplementary figure 5. Pathway-gene mutation heatmaps showing within pathway gene mutual exclusivity.** Heatmap of mutated genes in Transcriptional regulation by RUNX3 pathway in TARGET B-ALL tumors (#tumors: 162) showing the mutual exclusivity of *KRAS* & *TP53*.

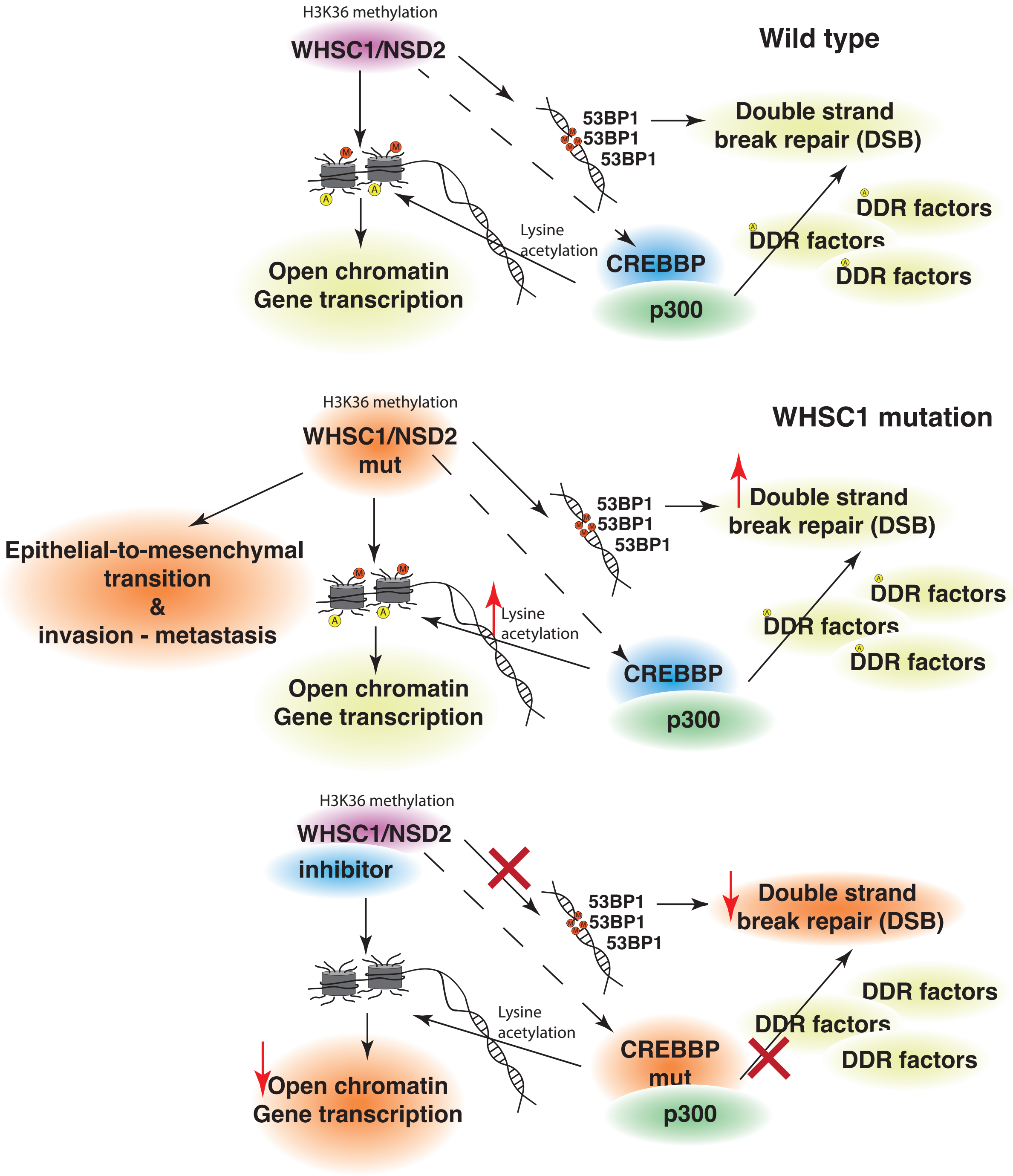


**Supplementary figure 6. Biological interpretation of candidate synthetic lethality between *WHSC1* and *CREBBP*.** Top: Role of *WHSC1*, *CREBBP* in DNA repair and transcription. *CREBBP/p300* acetylates DNA damage repair (DDR) factors, promoting DNA repair and regulating transcription modifying chromatin. *WHSC1* modifies chromatin by methylating histones at lysine 36 (H3K36 methylation) and methylates double strand breaks that are subsequently recognized by *53BP1* and promote DNA repair. Middle: In cancer, *WHSC1* mutations promote epithelial-to-mesenchymal transition and invasion. *WHSC1* promotes increased acetylation of chromatin by recruiting *CREBBP*. Bottom: Hypothesis of effect of inhibited *WHSC1* and mutated *CREBBP*. Inhibition of *WHSC1* and *CREBBP* mutation reduce open chromatin and impair DNA repair. Potentially this leads to accumulating unrepaired DNA to a point that could be synthetic sick or fatal for the cell.
