## Supplementary Table 5 for "A pathway-informed mutual exclusivity framework to detect genetic interactions in pediatric cancer"

### 1. Gene pair: TP53 & DROSHA in Wilms tumors

| | |
|---|---|
| **ME observed in articles:** | in Daub 2021 but in TARGET dataset (this case is derived from DKFZ dataset) |
| **Synthetic lethal observed in articles:** | - |
| **Main function:** | **TP53**: involved in DNA damage repair, cell cycle arrest [1]<br><br>**DROSHA**: DNA damage repair [2], microRNA biogenesis [3] |
| **Functionally related / Molecular interaction:** | When DNA damage occurs p53/p68 binds to DROSHA/DGCR8 (microprocessor complex) to affect miRNA biogenesis [4] |
| **Other:** | - |
| **This study's biological explanation – hypothesis:** | When both genes have a loss-of-function mutation likely have a great impact on repair mechanisms to a point that is intolerable and fatal. |

### 2. Gene pair: TP53 & DGCR8 in Wilms tumors

| | |
|---|---|
| **ME observed in articles:** | - |
| **Synthetic lethal observed in articles:** | - |
| **Main function:** | **TP53**: involved in DNA damage repair, cell cycle arrest (Speidel, 2015)<br><br>**DGCR8**: DNA damage repair (Hang et al., 2021), microRNA biogenesis (Ha & Kim, 2014) |
| **Functionally related / Molecular interaction:** | When DNA damage occurs p53/p68 binds to DROSHA/DGCR8 (microprocessor complex) to affect miRNA biogenesis [4] |
| **Other:** | - |
| **This study's biological explanation – hypothesis:** | When both genes have a loss-of-function mutation likely have a great impact on repair mechanisms to a point that is intolerable and fatal. |

### 3. Gene pair: KMT2D & NRAS in B-ALL

| | |
|---|---|
| **ME observed in articles:** | - |
| **Synthetic lethal observed in articles:** | - |

| | |
|---|---|
| **Main function:** | **KMT2D**: H3K4 methyltransferase, involved in several signaling & developmental pathways [6–9]<br><br>**NRAS**: GTPase that regulates MAPK and PI3K pathways. Promotes proliferation, survival, differentiation & cell-cell, cell-extracellular matrix interactions [10–12] |
| **Functionally related / Molecular interaction:** | KMT2D loss leads to activation of RTK-RAS signaling in lung squamous cell carcinoma, and hypersensitivity to RTK-RAS inhibition [13] |
| **Other:** | KMT (MLL) rearranged genes are associated with specific pediatric B-ALL subtypes [14]<br><br>Ras mutations drive leukemia via KMT2A-PLK1 axis [15] |
| **This study's biological explanation – hypothesis:** | Likely pathway epistasis. |

4. Gene pair: UBA2 & NRAS in B-ALL

| | |
|---|---|
| **ME observed in articles:** | **-** |
| **Synthetic lethal observed in articles:** | **-** |
| **Main function:** | **UBA2**: ubiquitin-like modifier-activating enzyme 2 forms a complex with SAE1 to activate SUMO proteins, stabilizing proteins to avoid degradation e.g. Rad23 [16], role in double-strand break resection [17]<br><br>**NRAS**: GTPase that regulates MAPK and PI3K pathways. Promotes proliferation, survival, differentiation & cell-cell, cell-extracellular matrix interactions [10–12] |
| **Functionally related / Molecular interaction:** | SUMO pathway can regulate Ras/MAPK [18] |
| **Other:** | - |
| **This study's biological explanation – hypothesis:** | Likely pathway epistasis. |

5. Gene pair: KRAS & KMT2B in B-ALL

| | |
|---|---|
| **ME observed in articles:** | **-** |
| **Synthetic lethal observed in articles:** | **-** |

| | |
|---|---|
| **Main function:** | **KRAS**: GTPase, important role in regulating proliferation, signaling in MAPK and PI3K pathway [19,20]<br><br>**KMT2B**: histone methyltransferase, involved in making genome accessible, transcription and gene expression, involved in cell cycle progression [21,22] |
| **Functionally related / Molecular interaction:** | - |
| **Other:** | Ras mutations drive leukemia via KMT2A-PLK1 axis [15] |
| **This study's biological explanation – hypothesis:** | Insufficient findings. |

### 6. Gene pair: UBA2 & KRAS in B-ALL

| | |
|---|---|
| **ME observed in articles:** | **-** |
| **Synthetic lethal observed in articles:** | **-** |
| **Main function:** | **UBA2**: ubiquitin-like modifier-activating enzyme 2 forms a complex with SAE1 to activate SUMO proteins, stabilizing proteins to avoid degradation e.g. Rad23 [16], role in double-strand break resection [17]<br><br>**KRAS**: GTPase, important role in regulating proliferation, signaling in MAPK and PI3K pathway [19,20] |
| **Functionally related / Molecular interaction:** | SUMO pathway can regulate Ras/MAPK [18] |
| **Other:** | - |
| **This study's biological explanation – hypothesis:** | Likely pathway epistasis. |

### 7. Gene pair: CTCF & KRAS in B-ALL

| | |
|---|---|
| **ME observed in articles:** | **-** |
| **Synthetic lethal observed in articles:** | **-** |
| **Main function:** | **CTCF**: mediates intrachromosomal and intrachromosomal interactions, regulates alternative mRNA splicing, enhancer-promoter interactions, recombination etc. [23] |

| | |
|---|---|
| | **KRAS**: GTPase, important role in regulating proliferation, signaling in MAPK and PI3K pathway [19,20] |
| **Functionally related / Molecular interaction:** | - |
| **Other:** | - |
| **This study's biological explanation – hypothesis:** | Insufficient information. |

### 8. Gene pair: FLT3 & KMT2D in B-ALL

| | |
|---|---|
| **ME observed in articles:** | - |
| **Synthetic lethal observed in articles:** | **-** |
| **Main function:** | **FLT3**: involved in accumulation of DNA damage in AMLs usually due to FLT3/ITD mutations [24,25], though not reported specifically in the context of ALL or B-ALL. Survival, proliferation and differentiation of hematopoietic cells [26]<br><br>**KMT2D**: H3K4 methyltransferase, involved in several signaling & developmental pathways [6–9] |
| **Functionally related / Molecular interaction:** | **-** |
| **Other:** | **-** |
| **This study's biological explanation – hypothesis:** | Insufficient information. |

### 9. Gene pair: JAK2 & PAX5 in B-ALL

| | |
|---|---|
| **ME observed in articles:** | - |
| **Synthetic lethal observed in articles:** | **-** |
| **Main function:** | **JAK2**: involved in DNA damage only in AML or myeloproliferative neoplasms [27] Involved via JAK/STAT pathway in proliferation and survival, transcription activation of several mediators of cancer and inflammation [28,29]<br><br>**PAX5**: major role in b-cell development and maturation (b-cell commitment) [30] |

| | |
|---|---|
| **Functionally related / Molecular interaction:** | - |
| **Other:** | Gene fusion PAX5-JAK2 activate STAT5 and promote aggressive B-ALL [31] |
| **This study's biological explanation – hypothesis:** | Subtype [14] |

10. **Gene pair:** CREBBP & FLT3 in B-ALL

| | |
|---|---|
| **ME observed in articles:** | - |
| **Synthetic lethal observed in articles:** | - |
| **Main function:** | **CREBBP**: transcription co-activator together with p300 [32,33], histone & transcription factor acetylation for chromatin accessibility [34,35], direct regulator of DDR [36]<br><br>**FLT3**: involved in accumulation of DNA damage in AMLs usually due to FLT3/ITD mutations [24,25], though not reported specifically in the context of ALL or B-ALL. Survival, proliferation and differentiation of hematopoietic cells [26] |
| **Functionally related / Molecular interaction:** | - |
| **Other:** | - |
| **This study's biological explanation – hypothesis:** | Insufficient information. |

11. **Gene pair:** JAK2 & WHSC1 in B-ALL

| | |
|---|---|
| **ME observed in articles:** | - |
| **Synthetic lethal observed in articles:** | - |
| **Main function:** | **JAK2**: involved in DNA damage only in AML or myeloproliferative neoplasms [27],<br>Involved via JAK/STAT pathway in proliferation and survival, transcription activation of several mediators of cancer and inflammation[28,29]<br><br>**WHSC1**(NSD2): histone methyltransferase, plays a role in initiating NHEJ by H3K36 dimethylation [37–39] |

| | |
|---|---|
| **Functionally related / Molecular interaction:** | - |
| **Other:** | - |
| **This study's biological explanation – hypothesis:** | Insufficient information. |

#### 12. **Gene pair:** CREBBP & JAK2 in B-ALL

| | |
|---|---|
| **ME observed in articles:** | - |
| **Synthetic lethal observed in articles:** | - |
| **Main function:** | **CREBBP**: transcription co-activator together with p300 [32,33], histone & transcription factor acetylation for chromatin accessibility [34,35], direct regulator of DDR [36]<br><br>**JAK2**: involved in DNA damage only in AML or myeloproliferative neoplasms [27], involved via JAK/STAT pathway in proliferation and survival, transcription activation of several mediators of cancer and inflammation [28,29] |
| **Functionally related / Molecular interaction:** | - |
| **Other:** | - |
| **This study's biological explanation – hypothesis:** | Insufficient information. |

#### 13. Gene pair: SYNE1 & PTPN11 in B-ALL

| | |
|---|---|
| **ME observed in articles:** | - |
| **Synthetic lethal observed in articles:** | - |
| **Main function:** | **SYNE1**(or Nesprin-1): part of LINC complex (linker of the nucleskeleton and cytoskeleton) that mediates mechanical signals from cytoskeleton to nuclear lamina [40,41], plays a role in DNA damage repair pathways [41] |

| | |
|---|---|
| | **PTPN11**: regulates development, signaling pathways ; Ras/MAPK, PI3K/AKT [42] and overexpressed in leukemogenesis [43] |
| **Functionally related / Molecular interaction:** | - |
| **Other:** | - |
| **This study's biological explanation – hypothesis:** | Insufficient information. |

#### 14. Gene pair: ZEB2 & PTPN11 in B-ALL

| | |
|---|---|
| **ME observed in articles:** | - |
| **Synthetic lethal observed in articles:** | - |
| **Main function:** | **ZEB2**: transcription factor, significant for promoting epitherlial-to-mesenchymal transition (EMT) https://doi.org/10.1093/nar/gki965 , plays a role in development and differentiation, but also in cancer, metastasis, survival, apoptosis etc.[44], plays a role in hematopoeitic stem, progenitor cell diferentiation and lineage fidelity [45]<br><br>**PTPN11**: regulates development, signaling pathways ; Ras/MAPK, PI3K/AKT [42] and overexpressed in leukemogenesis [43] |
| **Functionally related / Molecular interaction:** | SHP-2 (PTPN11) upregulates ZEB1 to induce EMT [46] |
| **Other:** | ZEB1 plays a role in DNA damage response [47] |
| **This study's biological explanation – hypothesis:** | Likely pathway epistasis. |

#### 15. Gene pair: UBA2 & PTPN11 in B-ALL

| | |
|---|---|
| **ME observed in articles:** | - |
| **Synthetic lethal observed in articles:** | - |
| **Main function:** | **UBA2**: ubiquitin-like modifier-activating enzyme 2 forms a complex with SAE1 to activate SUMO proteins, stabilizing proteins to avoid degradation e.g. Rad23 [16], role in double-strand break resection [17] |

| | |
|---|---|
| | **PTPN11**: regulates development, signaling pathways ; Ras/MAPK, PI3K/AKT [42] and overexpressed in leukemogenesis [43] |
| **Functionally related / Molecular interaction:** | PTPN11 (Shp2) is a target of SUMOylation – SUMOylated Shp2 induces ERK activation [48] |
| **Other:** | - |
| **This study's biological explanation – hypothesis:** | Likely pathway epistasis. |

#### 16. **Gene pair:** CREBBP & WHSC1 in B-ALL

| | |
|---|---|
| **ME observed in articles:** | **-** |
| **Synthetic lethal observed in articles:** | **-** |
| **Main function:** | **CREBBP**: transcription co-activator together with p300 [32,33], histone & transcription factor acetylation for chromatin accessibility [34,35], direct regulator of DDR [36]<br><br>**WHSC1**(NSD2): histone methyltransferase, plays a role in initiating NHEJ by H3K36 dimethylation [37–39] |
| **Functionally related / Molecular interaction:** | NSD2 regulates NF-kappaBeta, NF-kappaBeta recuits its co-activator p300-CBP (CREBBP-EP300) to induce histone (hyper)acetylation [49] |
| **Other:** | CREBBP usually acquires a loss-of-function mutation, WHSC1 usually acquire a gain-of-function mutation |
| **This study's biological explanation – hypothesis:** | Likely synthetic lethal/sick if both of the genes have a loss-of-function mutation or if targeted with an inhibitor. |

#### 17. Gene pair: WHSC1 & JAK1 in B-ALL

| | |
|---|---|
| **ME observed in articles:** | **-** |
| **Synthetic lethal observed in articles:** | **-** |
| **Main function:** | **WHSC1**(NSD2): histone methyltransferase, plays a role in initiating NHEJ by H3K36 dimethylation [37–39]<br><br>**JAK1**: Involved via JAK/STAT pathway in proliferation and survival, transcription activation of several mediators of cancer and inflammation [28,29] |
| **Functionally related / Molecular interaction:** | **-** |

| | |
|---|---|
| **Other:** | - |
| **This study's biological explanation – hypothesis:** | Insufficient information. |

#### 18. Gene pair: TP53 & KMT2B in B-ALL

| | |
|---|---|
| **ME observed in articles:** | - |
| **Synthetic lethal observed in articles:** | - |
| **Main function:** | **TP53**: involved in DNA damage repair, cell cycle arrest [1]<br><br>**KMT2B**: histone methyltransferase, involved in making genome accessible, transcription and gene expression, involved in cell cycle progression [21,22] |
| **Functionally related / Molecular interaction:** | - |
| **Other:** | - |
| **This study's biological explanation – hypothesis:** | Insufficient information. |

#### 19. Gene pair: SYNE1 & TP53 in B-ALL

| | |
|---|---|
| **ME observed in articles:** | - |
| **Synthetic lethal observed in articles:** | - |
| **Main function:** | **TP53**: involved in DNA damage repair, cell cycle arrest [1]<br><br>**SYNE1**(or Nesprin-1): part of LINC complex (linker of the nucleskeleton and cytoskeleton) that mediates mechanical signals from cytoskeleton to nuclear lamina [40,41], plays a role in DNA damage repair pathways [41] |
| **Functionally related / Molecular interaction:** | - |
| **Other:** | - |
| **This study's biological explanation – hypothesis:** | Likely synthetic sick/lethal. TP53 loss-of-function can induce increase of DSBs. SYNE1 is instrumental for nuclear shape, centrosome localization and genome stability. Upon decreased level of SYNE1 DNA damage repair is decreased/affected. |

| | |
|---|---|
| | Upon loss of both likely higher DNA damage levels that are sick or fatal. |

### 20. **Gene pair:** JAK2 & SETD2 in B-ALL

| | |
|---|---|
| **ME observed in articles:** | **-** |
| **Synthetic lethal observed in articles:** | **-** |
| **Main function:** | **JAK2**: involved in DNA damage only in AML or myeloproliferative neoplasms [27], Involved via JAK/STAT pathway in proliferation and survival, transcription activation of several mediators of cancer and inflammation[28,29]<br><br>**SETD2**: tumor suppressor, involved in DNA mismatch repair (MMR) – loss of SETD2 results in increased spontaneous mutations [50], safeguards genomic integrity [51] |
| **Functionally related / Molecular interaction:** | **-** |
| **Other:** | **-** |
| **This study's biological explanation – hypothesis:** | Insufficient information. |

### 21. Gene pair: TP53 & NRAS in B-ALL

| | |
|---|---|
| **ME observed in articles:** | **-** |
| **Synthetic lethal observed in articles:** | **-** |
| **Main function:** | **TP53**: involved in DNA damage repair, cell cycle arrest [1]<br><br>**NRAS**: GTPase that regulates MAPK and PI3K pathways. Promotes proliferation, survival, differentiation & cell-cell, cell-extracellular matrix interactions [10–12] |
| **Functionally related / Molecular interaction:** | - |
| **Other:** | Near-hypodiploidy associated with Ras-signaling alterations (and others), Low-hypodiploidy associated with TP53 alterations (and others) [14] |
| **This study's biological explanation – hypothesis:** | Subtype |

### 22. Gene pair: FLT3 & KMT2B in B-ALL

| | |
|---|---|
| **ME observed in articles:** | - |
| **Synthetic lethal observed in articles:** | **-** |
| **Main function:** | **FLT3**: involved in accumulation of DNA damage in AMLs usually due to FLT3/ITD mutations [24,25], though not reported specifically in the context of ALL or B-ALL. Survival, proliferation and differentiation of hematopoietic cells [26]<br><br>**KMT2B**: histone methyltransferase, involved in making genome accessible, transcription and gene expression, involved in cell cycle progression [21,22] |
| **Functionally related / Molecular interaction:** | **-** |
| **Other:** | **-** |
| **This study's biological explanation – hypothesis:** | Insufficient information. |

### 23. Gene pair: JAK2 & KMT2B in B-ALL

| | |
|---|---|
| **ME observed in articles:** | - |
| **Synthetic lethal observed in articles:** | **-** |
| **Main function:** | **JAK2**: involved in DNA damage only in AML or myeloproliferative neoplasms [27], Involved via JAK/STAT pathway in proliferation and survival, transcription activation of several mediators of cancer and inflammation[28,29]<br><br>**KMT2B**: histone methyltransferase, involved in making genome accessible, transcription and gene expression, involved in cell cycle progression [21,22] |
| **Functionally related / Molecular interaction:** | **-** |
| **Other:** | **-** |
| **This study's biological explanation – hypothesis:** | Insufficient information. |

#### 24. Gene pair: JAK2 & IKZF1 in B-ALL

| | |
|---|---|
| **ME observed in articles:** | co-occurrence and with CDKN2A/B linked with poor prognosis in ALL [14,52] |
| **Synthetic lethal observed in articles:** | - |
| **Main function:** | **JAK2**: involved in DNA damage only in AML or myeloproliferative neoplasms [27], Involved via JAK/STAT pathway in proliferation and survival, transcription activation of several mediators of cancer and inflammation[28,29]<br><br>**IKZF1**: transcription factor, involved in in lymphoid differentiation [53,54], recruits chromatin remodelers and histone deacetylases[55] |
| **Functionally related / Molecular interaction:** | |
| **Other:** | CRFL2 rearrangement with JAK and IKZF1 mutations has poor prognosis in B-progenitor acute lymphoblastic leukemia [56] |
| **This study's biological explanation – hypothesis:** | Likely false positive. |

#### 25. Gene pair: JAK1 & IKZF1 in B-ALL

| | |
|---|---|
| **ME observed in articles:** | co-occurrence and with CDKN2A/B linked with poor prognosis in ALL [14,52] |
| **Synthetic lethal observed in articles:** | - |
| **Main function:** | **JAK1**: Involved via JAK/STAT pathway in proliferation and survival, transcription activation of several mediators of cancer and inflammation [28,29]<br><br>**IKZF1**: transcription factor, involved in in lymphoid differentiation [53,54], recruits chromatin remodelers and histone deacetylases[55] |
| **Functionally related / Molecular interaction:** | - |
| **Other:** | CRFL2 rearrangement with JAK and IKZF1 mutations has poor prognosis in B- |

| | |
|---|---|
| | progenitor acute lymphoblastic leukemia [56] |
| **This study's biological explanation – hypothesis:** | Likely false positive. |

### 26. Gene pair: CENPE & IKZF1 in B-ALL

| | |
|---|---|
| **ME observed in articles:** | - |
| **Synthetic lethal observed in articles:** | - |
| **Main function:** | **CENPE**: important for chromosome alignment during prometaphase and congression [58–60]<br><br>**IKZF1**: transcription factor, involved in in lymphoid differentiation [53,54], recruits chromatin remodelers and histone deacetylases[55] |
| **Functionally related / Molecular interaction:** | - |
| **Other:** | - |
| **This study's biological explanation – hypothesis:** | Insufficient information. |

### 27. Gene pair: SETD2 & RYR2 in B-ALL

| | |
|---|---|
| **ME observed in articles:** | - |
| **Synthetic lethal observed in articles:** | - |
| **Main function:** | **SETD2**: tumor suppressor, involved in DNA mismatch repair (MMR) – loss of SETD2 results in increased spontaneous mutations [50], safeguards genomic integrity [51]<br><br>**RYR2**: receptors that control calcium release from the sarcoplasmic reticulum, mostly found in cardiomyocytes [62] , calcium plays a role in signaling as a second messenger e.g. PI3K pathway [63] |
| **Functionally related / Molecular interaction:** | - |
| **Other:** | - |

| | |
|---|---|
| **This study's biological explanation – hypothesis:** | Insufficient information. |

### 28. Gene pair: JAK1 & SETD2 in B-ALL

| | |
|---|---|
| **ME observed in articles:** | **-** |
| **Synthetic lethal observed in articles:** | **-** |
| **Main function:** | **JAK1**: Involved via JAK/STAT pathway in proliferation and survival, transcription activation of several mediators of cancer and inflammation [28,29]<br><br>**SETD2**: tumor suppressor, involved in DNA mismatch repair (MMR) – loss of SETD2 results in increased spontaneous mutations [50], safeguards genomic integrity [51] |
| **Functionally related / Molecular interaction:** | **-** |
| **Other:** | **-** |
| **This study's biological explanation – hypothesis:** | Insufficient information. |

### 29. Gene pair: KRAS & NPM1 in AML

| | |
|---|---|
| **ME observed in articles:** | **-** |
| **Synthetic lethal observed in articles:** | **-** |
| **Main function:** | **KRAS**: GTPase, important role in regulating proliferation, signaling in MAPK and PI3K pathway [19,20]<br><br>**NPM1**: involved in ribosome biogenesis and maturation [65–67], chaperone role for stability and localization of tumor suppressor proteins [68] , involved in DNA repair mechanisms [69] |
| **Functionally related / Molecular interaction:** | **-** |
| **Other:** | **-** |
| **This study's biological explanation – hypothesis:** | Insufficient information. |

### 30. Gene pair: NPM1 & NRAS in AML

| | |
|---|---|
| **ME observed in articles:** | - |
| **Synthetic lethal observed in articles:** | - |
| **Main function:** | **NRAS**: GTPase that regulates MAPK and PI3K pathways. Promotes proliferation, survival, differentiation & cell-cell, cell-extracellular matrix interactions [10–12]<br><br>**NPM1**: involved in ribosome biogenesis and maturation [65–67], chaperone role for stability and localization of tumor suppressor proteins [68] , involved in DNA repair mechanisms [69] |
| **Functionally related / Molecular interaction:** | - |
| **Other:** | - |
| **This study's biological explanation – hypothesis:** | Insufficient information. |

### 31. Gene pair: TP53 & PTCH1 in MB-SHH

| | |
|---|---|
| **ME observed in articles:** | yes [70,71] |
| **Synthetic lethal observed in articles:** | - |
| **Main function:** | **TP53**: involved in DNA damage repair, cell cycle arrest [1]<br><br>**PTCH1**: tumor suppressor, part of Sonic Hedgehog pathway (SHH), inhibits SMO oncogenic signaling [72] |
| **Functionally related / Molecular interaction:** | yes, in cancer SHH pathway, via for example Smo with an activating mutation, inhibits accumulation of TP53, therefore promoting cell proliferation [73] |
| **Other:** | - |
| **This study's biological explanation – hypothesis:** | Pathway epistasis. |

### 32. Gene pair: SMO & TP53 in MB-SHH

| | |
|---|---|
| **ME observed in articles:** | - |

| | |
|---|---|
| **Synthetic lethal observed in articles:** | - |
| **Main function:** | **SMO**: part of Sonic Hedgehog pathway (SHH), inhibited by PTCH1 [72]<br><br>**TP53**: involved in DNA damage repair, cell cycle arrest [1] |
| **Functionally related / Molecular interaction:** | yes, in cancer SHH pathway, via for example Smo with an activating mutation, inhibits accumulation of TP53, therefore promoting cell proliferation [73] |
| **Other:** | - |
| **This study's biological explanation – hypothesis:** | Pathway epistasis. |

### 33. Gene pair: CACNA1A & SLC45A3 in NBL

**This study's biological explanation – hypothesis:**

False positive based on oncoprint.

### 34. Gene pair: FBXW7 & PIK3R1 in T-ALL

| | |
|---|---|
| **ME observed in articles:** | - |
| **Synthetic lethal observed in articles:** | - |
| **Main function:** | **FBXW7**: important for SCF complex to identify its substrate and ubiquitinate it, several of its targets regulate cell proliferation & cycle, for example cyclin E, c-Myc, mTOR etc. [74]. Dysfunctional FBXW7 promotes cell division and enhances chromosomal instability. [75]<br><br>**PIK3R1**: subunit of PI3K [76], PI3K activating AKT, which AKT downstream controls cell proliferation [77] |
| **Functionally related / Molecular interaction:** | - |
| **Other:** | - |
| **This study's biological explanation – hypothesis:** | Insufficient information. |

#### 35. Gene pair: DNM2 & LEF1 in T-ALL

| | |
|---|---|
| **ME observed in articles:** | - |
| **Synthetic lethal observed in articles:** | - |
| **Main function:** | **DNM2**: oncogene, dynamin 2 plays a role in endocytosis and intracellular membrane trafficking [78], regulates T-cell activation [79], involved in migration, invasion, metastasis & cell proliferation and survival pathways [80]<br><br>**LEF1**: oncogene, T-cell and B-cell enhancer transcription factor [81], involved in Wnt pathway [82] in epithelial-mesenchymal transition (EMT) |
| | - |
| **Other:** | - |
| **This study's biological explanation – hypothesis:** | Insufficient information. |

#### 36. Gene pair: LEF1 & IL7R in T-ALL

| | |
|---|---|
| **ME observed in articles:** | - |
| **Synthetic lethal observed in articles:** | - |
| **Main function:** | **LEF1**: oncogene, T-cell and B-cell enhancer transcription factor [81], involved in Wnt pathway [82] in epithelial-mesenchymal transition (EMT)<br><br>**IL7R**: oncogene, binds IL-7, involved in the JAK-STAT pathway [83] role in V(D)J recombination in lymphocyte development [84,85], important for T-cell development [86] |
| **Functionally related / Molecular interaction:** | - |
| **Other:** | - |
| **This study's biological explanation – hypothesis:** | Insufficient information. |

#### 37. Gene pair: RUNX1 & JAK1 in T-ALL

| | |
|---|---|
| **CO observed in articles:** | **-** |
| **Cooperation observed in articles:** | **-** |
| **Main function:** | **RUNX1**: tumor suppressor, transcription factor major regulator in hematopoietic differentiation, interacts with transcription complexes, interacts with histone modifiers to activate or repress transcription [87–89]<br><br>**JAK1**: Involved via JAK/STAT pathway in proliferation and survival, transcription activation of several mediators of cancer and inflammation [28,29] |
| **Functionally related / Molecular interaction:** | RUNX1 represses JAK/STAT signaling and sensitizes leukemic cells to JAK inhibition https://pubmed.ncbi.nlm.nih.gov/37581927/ |
| **Other:** | **-** |
| **This study's biological explanation – hypothesis:** | False positive. |
